# Progressive neural organization across the IFG and pMTG tracks gesture–speech information-theoretic structure

**DOI:** 10.1101/2022.11.23.517759

**Authors:** Wanying Zhao, Zhouyi Li, Xiang Li, Yi Du

**Affiliations:** State Key Laboratory of Cognitive Science and Mental Health, Institute of Psychology, Chinese Academy of Sciences, Beijing, China 100101; Department of Psychology, University of Chinese Academy of Sciences, Beijing, China 100049

**Author notes:** Corresponding author: Dr. Yi Du, 16 Lincui Road, Chaoyang district, Beijing, China 100101 Dr. Wanying Zhao, 16 Lincui Road, Chaoyang District, Beijing, China 100101.

**Keywords:** gesture-speech integration, pMTG-IFG network, information theory, multisensory, semantic

## Abstract

Semantic representation emerges from distributed multisensory modalities, yet a comprehensive understanding of the functional changing pattern within convergence zones or hubs integrating multisensory semantic information remains elusive. In this study, employing information-theoretic metrics, we quantified gesture and speech information, alongside their interaction, utilizing entropy and mutual information (MI). Neural implementation of these information-theoretic representations was characterized across three complementary experiments using high-definition transcranial direct current stimulation (HD-tDCS), functional magnetic resonance imaging (fMRI), and electroencephalography (EEG). HD-tDCS demonstrated a causal contribution of the left inferior frontal gyrus (IFG) and posterior middle temporal gyrus (pMTG) to neural representations associated with gesture-speech MI. fMRI revealed that MI is represented across IFG and pMTG through distinct mechanisms: IFG preserves the relational structure predicted by MI, whereas pMTG supports discrimination of different levels of MI. EEG further showed that entropy primarily reflects transient modality-specific dynamics, whereas MI emerges during the N400-associated integration period. Cross-modal EEG–fMRI analyses demonstrated convergent MI-related neural representations with complementary temporal and spatial signatures. Together, these findings reveal a progressive organization of multisensory semantic integration, in which neural representations systematically track stimulus-level information-theoretic structure, with modality-specific uncertainty and cross-modal shared information expressed at different neural scales. This work establishes an information-theoretic framework that links the statistical structure of sensory information with its neural organization, providing new insights into how distributed inputs are coordinated to form coherent semantic representations.

## Introduction

Semantic representation, distinguished by its cohesive conceptual nature, emerges from distributed modality-specific regions. Consensus acknowledges the presence of ‘convergence zones’ within the temporal and inferior parietal areas^1^, or the ‘semantic hub’ located in the anterior temporal lobe^2^, pivotal for integrating, converging, or distilling multimodal inputs. Contemporary theories frame the semantic processing as a dynamic sequence of neural states^3^, shaped by systems that are finely tuned to the statistical regularities inherent in sensory inputs^4^. These regularities enable the brain to evaluate, weight, and integrate multisensory information, optimizing the reliability of individual sensory signals^5^. However, sensory inputs available to the brain are often incomplete and uncertain, necessitating adaptive neural adjustments to resolve these ambiguities^6^. In this context, neuronal activity is thought to be linked to the probability density of sensory information, with higher levels of uncertainty resulting in the engagement of a broader population of neurons, thereby reflecting the brain’s adaptive capacity to handle diverse possible interpretations^7,8^. Although the role of ‘convergence zones’ and ‘semantic hubs’ in integrating multimodal inputs is well established, the precise functional patterns of neural activity in response to the distribution of unified multisensory information—along with the influence of unisensory signals —remain poorly understood.

To this end, we developed an analytic approach to directly probe the cortical engagement during multisensory gesture-speech semantic integration. Even though gestures convey information in a global-synthetic way, while speech conveys information in a linear segmented way, there exists a bidirectional semantic influence between the two modalities^9,10^. Gesture is regarded as ‘part of language’^11^ or functional equivalents of lexical units that alternate and integrate with speech into a ‘single unification space’ to convey a coherent meaning^12–14^. Empirical studies have investigated the semantic integration between gesture and speech by manipulating their semantic relationship^15–18^ and revealed a mutual interaction between them^19–21^ as reflected by the N400 latency and amplitude^14^ as well as common neural underpinnings in the left inferior frontal gyrus (IFG) and posterior middle temporal gyrus (pMTG)^15,22,23^.

Building on these insights, the present study quantified the amount of information from both sources and their interaction adopting the information-theoretic complexity metrics of *entropy* and *mutual information* (MI). Unisensory Entropy measures the disorder or randomness of information and serves as an index of the uncertainty in modality-specific representations of gesture or speech activated by an event^24^. MI assesses share information between modalities^25^, indicating multisensory convergence and acting as an index of gesture-speech integration (**Figure 1A**).

**Figure 1.**
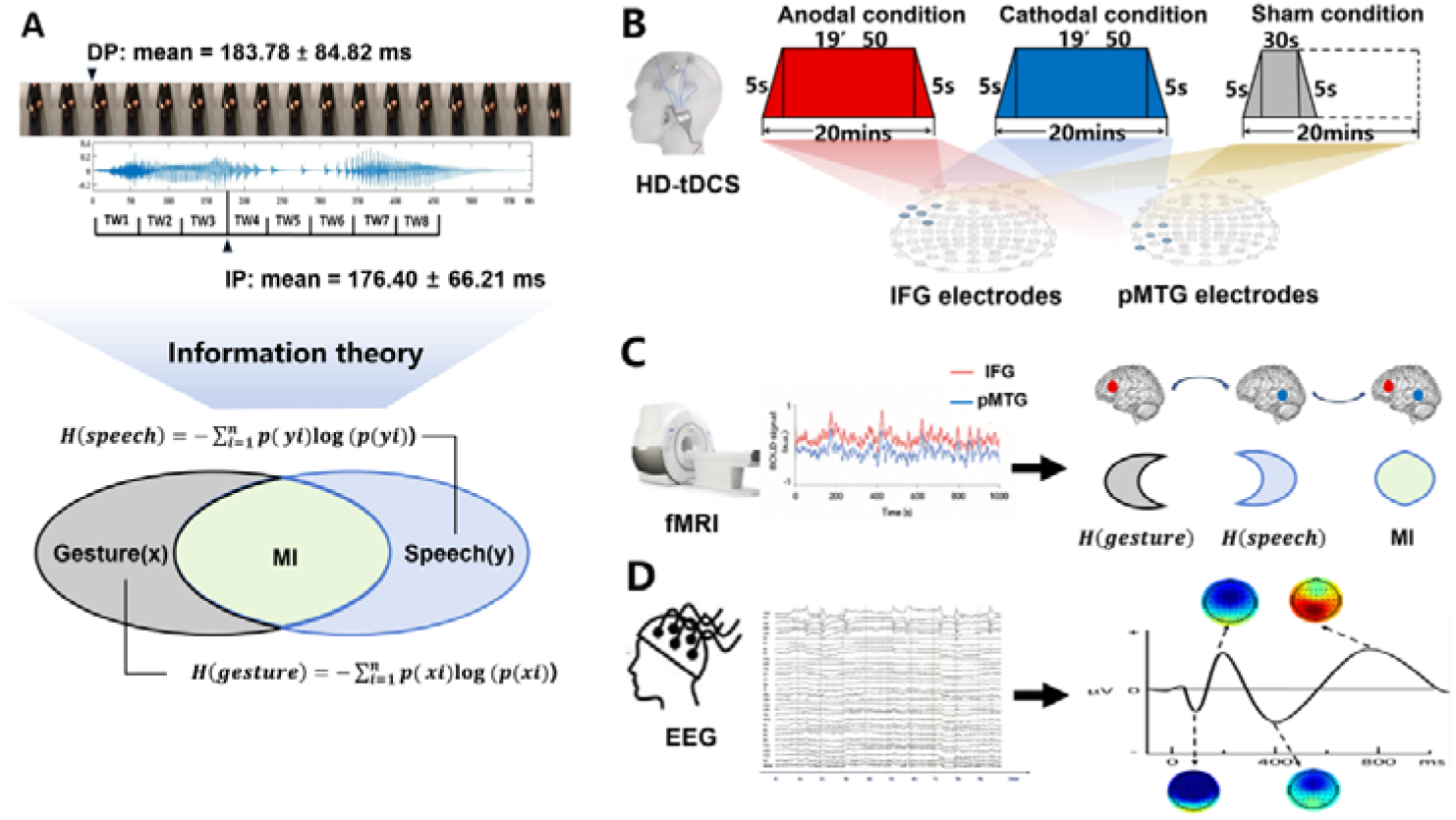
Experimental design, and stimulus characteristics. **(A) Experimental stimuli.** Twenty gestures were paired with 20 relevant speech stimuli. Two separate Pre-tests were executed to define the minimal length of each gesture and speech required for semantic identification, namely, the discrimination point (DP) of gesture and the identification point (IP) of speech. Overall, a mean of 183.78 ms (SD = 84.82) was found for the DP of gestures and the IP of speech was 176.40 ms (SD = 66.21). The onset of speech was set at the gesture DP. Responses for each item were assessed utilizing information-theoretic complexity metrics to quantify the information content of both gesture and speech during integration, employing entropy and MI. **(B) Procedure of Experiment 1.** HD-tDCS, including Anodal, Cathodal, or Sham conditions, was administered to the IFG or pMTG using a 4 * 1 ring-based electrode montage. Electrode F7 targeted the IFG, with return electrodes placed on AF7, FC5, F9, and FT9. For pMTG stimulation, TP7 was targeted, with return electrodes positioned on C5, P5, T9, and P9. Sessions lasted 20 minutes, with a 5-second fade-in and fade-out, while the Sham condition involved only 30 seconds of stimulation. **(C) Procedure of Experiment 2.** Blood-oxygen-level-dependent (BOLD) signals were acquired using functional magnetic resonance imaging (fMRI) while participants viewed semantically congruent gesture–speech stimuli. Region-specific neural representations were extracted from the left IFG and pMTG, and were related to stimulus-level information-theoretic metrics, including gesture entropy, speech entropy, and MI. Dissociable spatial representations of different information-theoretic metrics across the IFG–pMTG network were hypothesized. **(D) Procedure of Experiment 3.** Semantically congruent gesture-speech pairs were presented randomly with Electroencephalogram (EEG) recorded simultaneously. Epochs were time locked to the onset of speech and lasted for 1000 ms. A 200 ms pre-stimulus baseline correction was applied before the onset of gesture stoke. Various elicited components were hypothesized.

To investigate the neural mechanisms underlying gesture-speech integration, we conducted three experiments to assess how neural activity correlates with distributed multisensory integration, quantified using information-theoretic measures of MI. Additionally, we examined the contributions of unisensory signals in this process, quantified through unisensory entropy. We hypothesized that multisensory integration is progressively organized by information-theoretic structure, such that modality-specific and cross-modal information representations are differentially expressed across neural systems and processing stages. **Experiment 1** employed high-definition transcranial direct current stimulation (HD-tDCS) to establish the causal contribution of IFG and pMTG to neural representations associated with information-theoretic properties of gesture–speech integration. Anodal, Cathodal and Sham stimulation were applied to either the IFG or the pMTG (**Figure 1B**). HD-tDCS induces membrane depolarization with anodal stimulation and membrane hyperpolarization with cathodal stimulation^26^, thereby increasing or decreasing cortical excitability in the targeted brain area, respectively. By comparing stimulation-induced changes in behavioral performance across stimulation conditions, this experiment examined whether perturbing the IFG and/or pMTG alters sensitivity to the information structure of multisensory stimuli.

**Experiment 2** used functional magnetic resonance imaging (fMRI) to characterize the spatial organization of information-theoretic representations across IFG and pMTG semantic network (**Figure 1C**). Building on the causal contribution identified in Experiment 1, this experiment examined how information-theoretic properties of gesture–speech stimuli are represented within and coordinated across regions supporting multisensory semantic integration. Through complementary representational analyses, including representational similarity analysis (RSA), multivoxel pattern analysis (MVPA), and representational connectivity analysis, we investigated whether different information-theoretic metrics exhibit distinct spatial representational profiles and whether IFG and pMTG share or differentiate stimulus-level information structures.

**Experiment 3** complemented these investigations by focusing on the temporal dynamics of neural responses during semantic processing, leveraging high-temporal event-related potentials (ERPs). This experiment investigated how distinct information contributors unfold over time and are expressed in temporally distinct neural responses during semantic processing (**Figure 1D**). These temporal dynamics were characterized through ERP components included the early sensory effects as P1 and N1–P2^27,28^, the N400 semantic conflict effect^14,28,29^, and the late positive component (LPC) reconstruction effect^30,31^. By combining ERP analyses with representational geometry and multivariate neural pattern analyses within IFG- and pMTG-approximating electrode clusters, we investigated the temporal emergence of entropy- and MI-related representations across processing stages. Integrating these temporal dynamics with the causal contribution identified in Experiments 1 and the spatial organization revealed by fMRI in Experiment 2, Experiment 3 aimed to determine how modality-specific and cross-modal convergence are dynamically expressed during multisensory semantic integration.

## Results

### Experiment 1

#### Modulation of left pMTG and IFG engagement by gradual changes in gesture-speech semantic information

In the IFG, one-way ANOVA examining the effects of three tDCS conditions (Anodal, Cathodal, or Sham) on semantic congruency (RT (semantic incongruent) – RT (semantic congruent)) demonstrated a significant main effect of stimulation condition (*F*(2, 75) = 3.673, *p* = 0.030, ηp2 = 0.089). Post hoc paired t-tests indicated a significantly reduced semantic congruency effect between the Cathodal condition and the Sham condition (*t*(26) = −3.296, *p* = 0.003, 95% CI = [−11.488, 4.896]) (**Figure 3A left**). Subsequent Pearson correlation analysis revealed that the reduced semantic congruency effect was progressively associated with the MI, evidenced by a significant correlation between the Cathodal-tDCS effect (Cathodal-tDCS minus Sham-tDCS) and MI (*r* = −0.595, *p* = 0.007, 95% CI = [−0.995, −0.195]) (**Figure 3B**). Additionally, a similar correlation was observed between the Cathodal-tDCS effect and the total response number (*r* = −0.543, *p* = 0.016, 95% CI = [−0.961, −0.125]).

**Figure 2.**
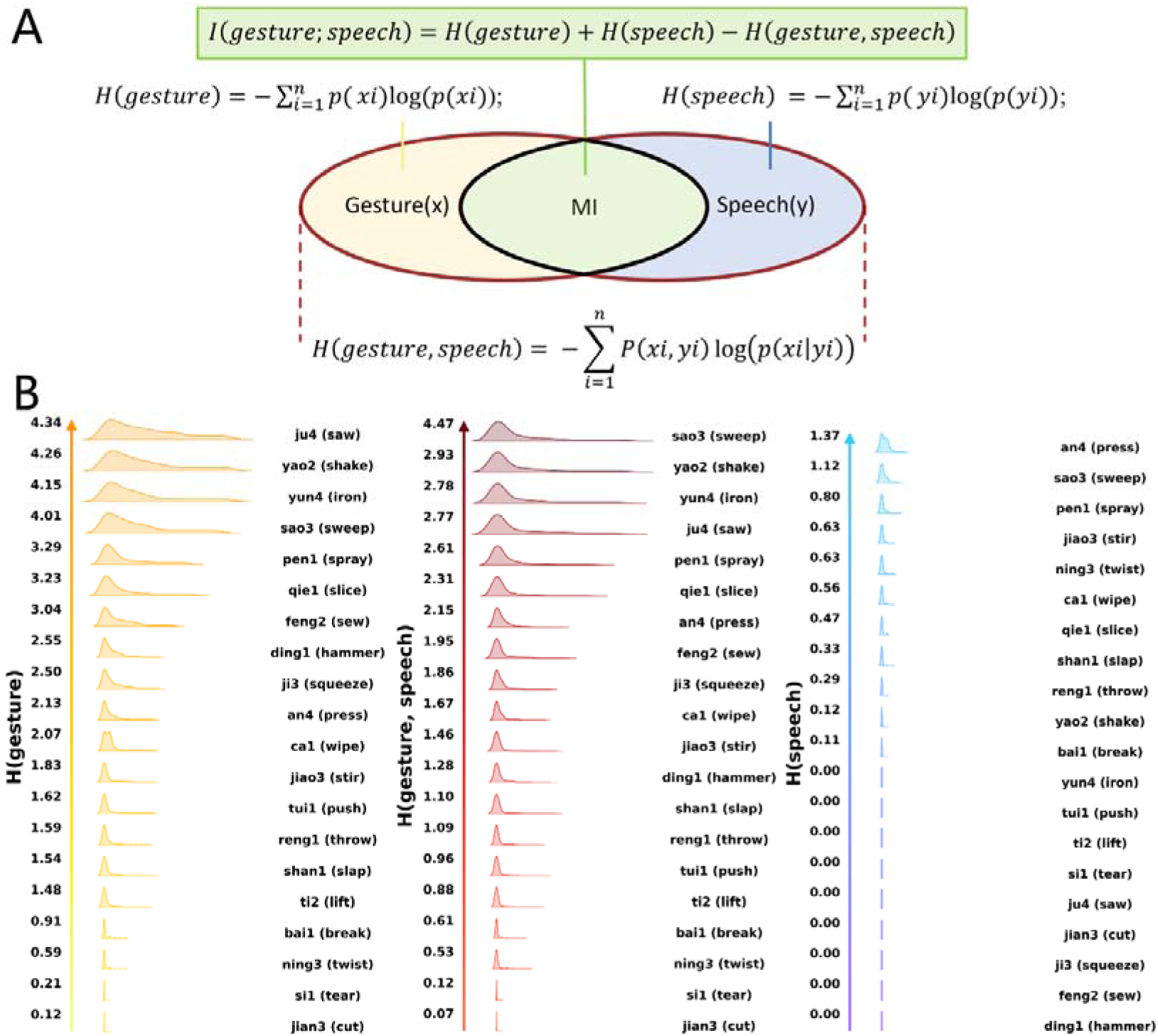
Quantification formulas (A) and distributions of each stimulus in Shannon’s entropy (B). Two separate pre-tests (N = 30) were conducted to assign a single verb for describing each of the isolated 20 gestures and 20 speech items. Responses provided for each item were transformed into Shannon’s entropy using a relative quantification formula. Gesture (**B left**) and speech (**B right**) entropy quantify the randomness of gestural or speech information, representing the uncertainty of probabilistic representation activated when a specific stimulus occurs. Joint entropy (**B middle**) captures the widespread nature of the two sources of information combined. Mutual information (MI) was calculated as the difference between joint entropy with gesture entropy and speech entropy combined (**A**), thereby capturing the overlap of gesture and speech and representing semantic integration.

**Figure 3.**
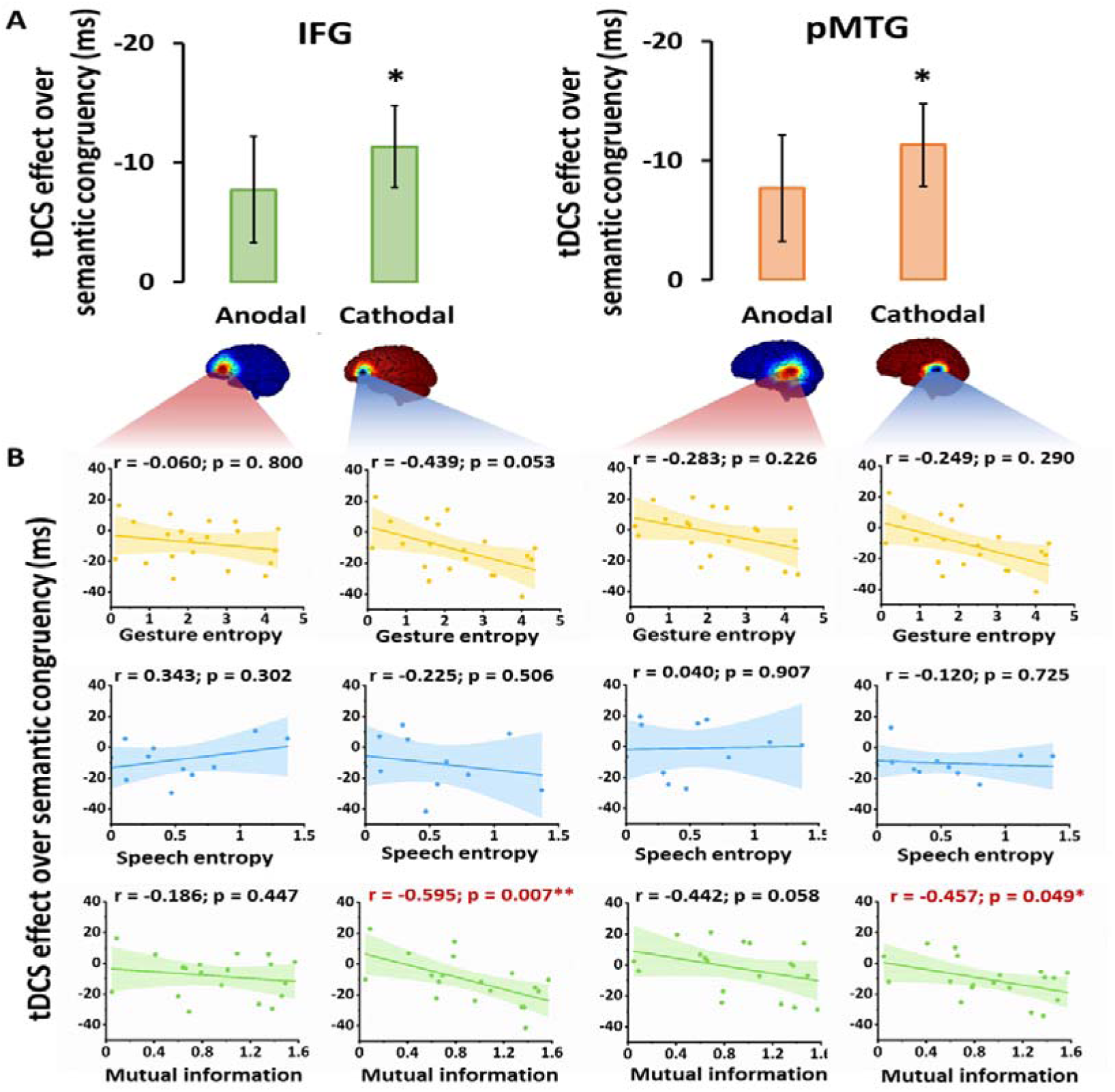
tDCS effect over semantic congruency. **(A)** tDCS effect was defined as active-tDCS minus sham-tDCS. The semantic congruency effect was calculated as the reaction time (RT) difference between semantically incongruent and semantically congruent pairs (Rt(incongruent) - Rt(congruent)). **(B)** Correlations of the tDCS effect over the semantic congruency effect with three information models (gesture entropy, speech entropy and MI) are displayed with best-fitting regression lines. Significant correlations are marked in red. * *p* < 0.05, ** *p* < 0.01 after FDR correction.

However, partial correlation analysis, controlling for the total response number, revealed that the initially significant correlation between the Cathodal-tDCS effect and MI was no longer significant (*r* = −0.303, *p* = 0.222, 95% CI = [−0.770, 0.164]). This suggests that the observed relationship between Cathodal-tDCS and MI may be confounded by semantic control difficulty, as reflected by the total number of responses. Specifically, the reduced activity in the IFG under Cathodal-tDCS may be driven by variations in the difficulty of semantic control rather than a direct modulation of MI.

In the pMTG, a one-way ANOVA assessing the effects of three tDCS conditions on semantic congruency also revealed a significant main effect of stimulation condition (*F*(2, 75) = 3.250, *p* = 0.044, ηp2 = 0.080). Subsequent paired t-tests identified a significantly reduced semantic congruency effect between the Cathodal condition and the Sham condition (*t*(25) = - 2.740, *p* = 0.011, 95% CI = [−11.915, 6.435]) (**Figure 3A right**). Moreover, a significant correlation was observed between the Cathodal-tDCS effect and MI (*r* = −0.457, *p* = 0.049, 95% CI = [−0.900, −0.014]) (**Figure 3B**).

Importantly, the reduced activity in the pMTG under Cathodal-tDCS was not influenced by the total response number, as indicated by the non-significant correlation (*r* = −0.253, *p* = 0.295, 95% CI = [−0.735, 0.229]). This finding was further corroborated by the unchanged significance in the partial correlation between Cathodal-tDCS and MI, when controlling for the total response number (*r* = −0.472, *p* = 0.048, 95% CI = [−0.903, −0.041]).

RTs of congruent and incongruent trials of IFG and pMTG in each of the stimulation conditions were shown in **Supplementary Table 4.**

### Experiment 2

#### Information-theoretic representations are differentially organized within the IFG–pMTG network

To determine how information-theoretic properties of gesture–speech stimuli are represented within the cortical network supporting multimodal semantic integration, we performed complementary multivariate analyses within independently defined left IFG and pMTG regions of interest. These regions were independently defined from a whole-brain contrast comparing semantically incongruent and congruent trials (**Figure 4A left**), and analyses were restricted to correctly recognized semantically congruent trials to characterize successful multimodal integration.

**Figure 4.**
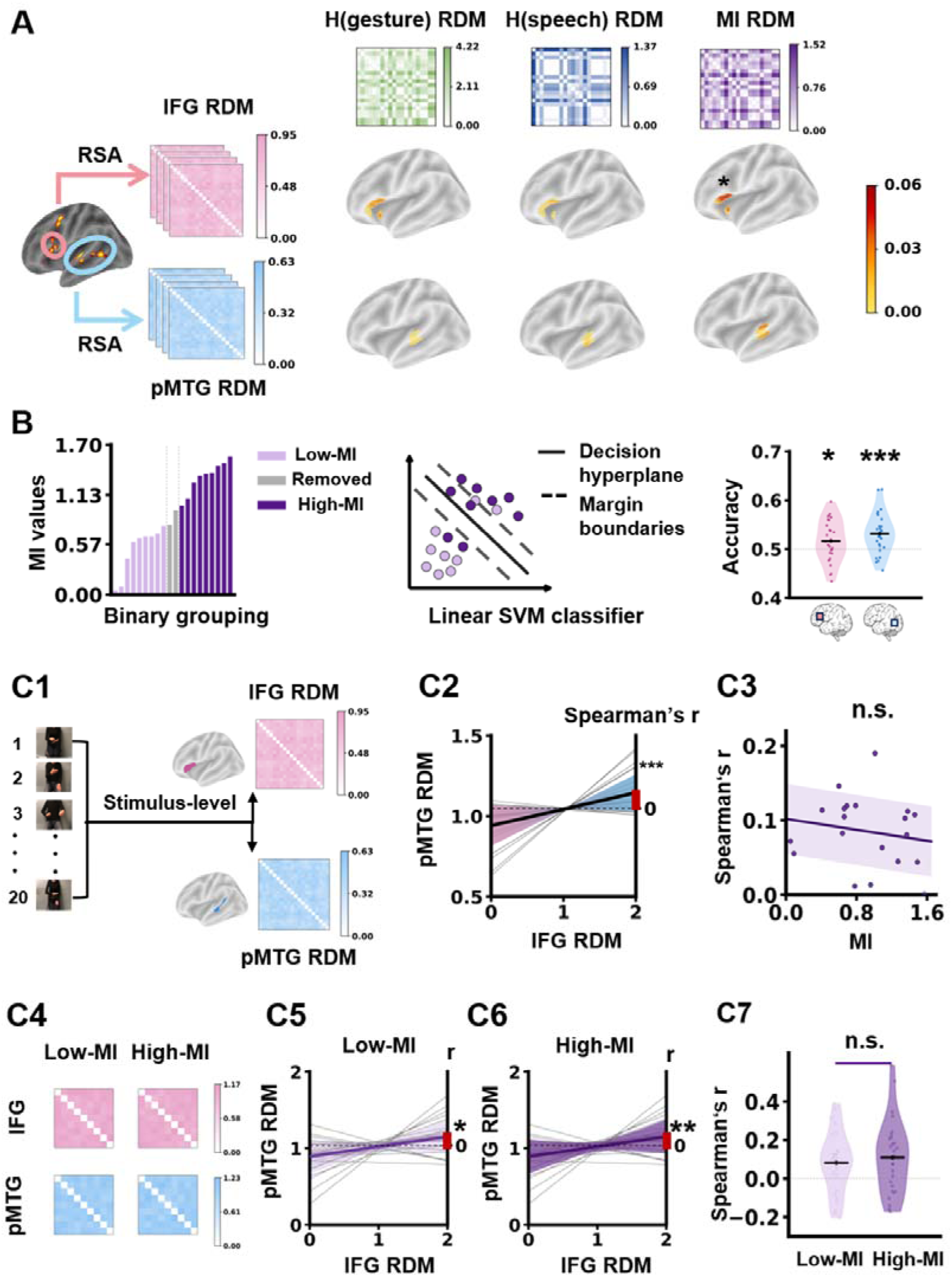
fMRI reveals distinct MI-related neural representations within the IFG–pMTG network. **(A)** Searchlight representational similarity analysis (RSA) within the left inferior frontal gyrus (IFG) and posterior middle temporal gyrus (pMTG) regions of interest, defined from a whole-brain semantic congruency contrast (incongruent > congruent trials). Neural representational dissimilarity matrices (RDMs) were compared with model RDMs derived from gesture entropy, speech entropy, and mutual information (MI). A significant MI-related representational structure was identified within the left IFG, whereas neither gesture entropy nor speech entropy showed significant correspondence with local neural geometry. **(B)** ROI-based multivoxel pattern analysis (MVPA) of MI-related information. Stimuli were ranked according to MI values and divided into low- and high-MI groups. Linear support vector machine (SVM) classifiers were trained using beta-series multivoxel patterns from the IFG and pMTG. Balanced decoding accuracy was significantly above chance in both regions. Violin plots show participant-level decoding accuracies; horizontal black lines indicate group means. **(C)** Representational connectivity analysis between IFG and pMTG. (**C1**) Schematic illustration of the representational connectivity analysis. Stimulus-level neural RDMs were generated separately for IFG and pMTG, and their correspondence was quantified using Spearman correlation. (**C2**) Overall IFG–pMTG representational connectivity across all stimuli. The mean Spearman correlation between IFG and pMTG RDMs was significantly greater than zero. The red bar indicates the group-level mean correlation coefficient tested against zero. (**C3**) Relationship between MI values and IFG–pMTG representational connectivity. Continuous MI variation did not significantly predict representational connectivity. (**C4**–**C6**) Representational connectivity for low- and high-MI stimulus subsets. IFG – pMTG RDM correlations were significant in both low-MI and high-MI conditions. (**C7**) Comparison of representational connectivity between low- and high-MI conditions. No significant difference was observed between conditions. * *p* < 0.05, ** *p* < 0.01, ** *p* < 0.001 after FDR correction.

We first examined whether local neural representational geometry was structured according to information-theoretic properties using searchlight representational similarity analysis (RSA). We then assessed whether regional multivoxel patterns contained discriminable information about high versus low information-theoretic states using multivoxel pattern analysis (MVPA). Finally, we quantified whether IFG and pMTG shared a common stimulus representation space using representational connectivity analysis.

#### IFG represents the representational geometry of MI

We first applied searchlight RSA within the IFG and pMTG regions to examine whether local neural representational geometry preserved the stimulus relationships predicted by information-theoretic models. RSA quantifies the correspondence between neural representational dissimilarity matrices (RDMs) and model RDMs derived from gesture entropy, speech entropy, and MI, thereby testing whether neural patterns preserve the relative structure of information-theoretic relationships across stimuli.

Within the left IFG, MI was the only information-theoretic dimension that significantly predicted local representational geometry. Specifically, a significant MI-sensitive cluster was identified within the IFG mask (cluster volume = 288 mm³; peak MNI coordinate = −44, 20, 4; cluster-level *p*FWE = 0.014). Neither gesture entropy nor speech entropy produced significant representational correspondence within this region. In contrast, no significant RSA clusters were detected for any information-theoretic dimension within the left pMTG (**Figure 4A right**). These results indicate that the left IFG contains local multivoxel patterns whose stimulus-level organization follows the degree of gesture–speech coupling captured by MI.

#### pMTG and IFG show dissociable representations of MI

To complement RSA, which characterizes whether neural representations preserve the continuous geometry of information-theoretic relationships, we performed ROI-based MVPA to examine whether these representations contained discriminable information about different information states. Stimuli were ranked according to their gesture entropy, speech entropy, or MI values, and linear classifiers were trained to distinguish stimuli with relatively high-versus low-values of each metric.

In the left pMTG, multivoxel patterns reliably discriminated high-versus low-MI stimuli (mean balanced accuracy = 0.531 ± 0.041; *t*(27) = 4.016, *p* < 0.001, Cohen’s d = 0.759) (**Figure 4B right**). In contrast, neither gesture entropy (accuracy = 0.514 ± 0.045; *t*(27) = 1.697, *p* = 0.101; d = 0.321) (**Supplementary figure 1A**) nor speech entropy (accuracy = 0.489 ± 0.048; *t*(27) = −1.262, *p* = 0.218; d = −0.238) (**Supplementary figure 1B**) could be reliably decoded.

In the left IFG, MI decoding also exceeded chance level, although with a smaller effect size (accuracy = 0.516 ± 0.042; *t*(27) = 2.063, *p* = 0.049, Cohen’s d = 0.390) (**Figure 4B right**). No significant decoding was observed for gesture entropy (accuracy = 0.504 ± 0.054; *t*(27) = 0.403, *p* = 0.690; d = 0.076) (**Supplementary figure 1A**) or speech entropy (accuracy = 0.494 ± 0.059; *t*(27) = −0.553, *p* = 0.585; d = −0.105) (**Supplementary figure 1B**).

Together, the RSA and MVPA revealed complementary but dissociable patterns of MI representation across the IFG–pMTG network. Whereas pMTG activity patterns contain stronger stimulus-discriminative information about cross-modal coupling states, IFG representations primarily preserve the relational structure underlying multimodal semantic organization. Thus, although both regions represent cross-modal information, they appear to contribute distinct aspects of MI-related processing.

#### IFG–pMTG connectivity does not account for MI-related functional differentiation

Although IFG and pMTG both contained MI-related representations, we next asked whether this functional differentiation was associated with MI-dependent changes in representational connectivity between the two regions. To address this question, we quantified representational connectivity by comparing stimulus-level neural RDMs between IFG and pMTG (**Figure 4C1**).

Across all stimuli, IFG and pMTG representational dissimilarity matrices showed a significantly correlated (mean Spearman *r* = 0.094 ± 0.102; *t*(27) = 4.787, *p* < 0.001) (**Figure 4C2**), indicating that the two regions exhibited partially similar stimulus-level representational organization during multimodal semantic processing.

We next tested whether the strength of IFG–pMTG representational connectivity varied as a function of cross-modal information. Stimuli were divided into high- and low-MI subsets, and representational connectivity was calculated separately for each subset (**Figure 4C4**).

Connectivity was significant for both high-MI (*r* = 0.110 ± 0.189; *t*(27) = 3.022, *p* = 0.005) (**Figure 4C6**) and low-MI subset (*r* = 0.082 ± 0.157; *t*(27) = 2.753, *p* = 0.010) (**Figure 4C5**). However, connectivity strength did not differ between high- and low-MI conditions (*t*(27) = 0.764, *p* = 0.452) (**Figure 4C7**).

Consistent with this result, continuous modulation analyses failed to reveal a significant relationship between MI values and IFG–pMTG representational connectivity (*t*(27) = −1.706, *p* = 0.099) (**Figure 4C3**). Comparable analyses using gesture entropy and speech entropy also failed to reveal systematic modulation of IFG–pMTG representational connectivity (all high–low comparisons and continuous correlations *p* > 0.05).

Together, these findings indicate that IFG and pMTG are functionally coordinated during multimodal semantic processing but that their distinct MI-related representational properties are not explained by MI-dependent modulation of inter-regional representational connectivity. Instead, the two regions appear to support complementary aspects of cross-modal information processing within the broader semantic network.

### Summary

Across complementary representational analyses, MI emerged as the information-theoretic dimension most consistently associated with neural organization within the IFG–pMTG network. Searchlight RSA revealed that IFG representations preserved the relational structure predicted by MI, whereas MVPA demonstrated that MI-related stimulus distinctions were more reliably decoded from pMTG patterns and, to a lesser extent, IFG patterns. Together, these findings indicate that IFG and pMTG contribute distinct aspects of MI representation during multimodal semantic integration, with IFG preserving the relational structure captured by MI and pMTG supporting discrimination of MI-defined stimulus differences.

Representational connectivity analysis further showed that IFG and pMTG exhibited significant stimulus-level representational correspondence, consistent with their participation in the same multimodal semantic network. However, the strength of this correspondence was not enhanced for high-MI stimuli and was not predicted by continuous MI values, indicating that the functional differentiation observed between IFG and pMTG does not arise from MI-dependent modulation of inter-regional representational coupling. Thus, although IFG and pMTG are anatomically and functionally connected in gesture-speech integration^22,23^, they appear to support complementary and relatively distinct aspects of MI-related representation.

### Experiment 3

#### EEG reveals distinct temporal dynamics of modality-specific entropy and MI during gesture-speech semantic integration

To determine how information-theoretic representations unfold over time, we recorded EEG responses from an independent cohort of participants and examined the temporal dynamics of gesture entropy, speech entropy, MI. While fMRI analyses identified MI as the reliable information-theoretic dimension represented within the IFG–pMTG network, EEG provided complementary temporal resolution to characterize when modality-specific entropy and cross-modal integration emerge during semantic processing.

Because the preceding experiments consistently implicated the left IFG and pMTG as critical regions supporting gesture–speech semantic integration, EEG analyses focused primarily on electrode clusters approximating these two regions. Complementary whole-scalp analyses were conducted to examine whether information-theoretic representations extended beyond these predefined regions and are reported in the Supplementary Information.

We first examined whether gesture entropy, speech entropy, and MI modulated stimulus-evoked neural responses using ROI-based ERP amplitude analyses. Subsequently, we applied time-resolved RSA and multivariate decoding to determine when information-theoretic structure and discriminative information emerged in neural representations.

#### Distinct temporal signatures of modality-specific entropy and MI

Linear mixed-effects models were used to assess whether stimulus-level ERP amplitudes were systematically modulated by gesture entropy, speech entropy, and MI, with participant variability modeled as a random effect.

In the left IFG region, MI showed a biphasic temporal profile. Higher MI values were associated with reduced positive activity during the early P1 window (β = −0.513, SE = 0.174, *t* = −2.944, *p*FDR = 0.014), followed by increased positivity during the N400 window (β = 0.409, SE = 0.163, *t* = 2.505, *p* = 0.025). No significant MI-related modulation was observed during N1/P2 or LPC windows. In contrast, MI did not significantly modulate ERP amplitudes within the pMTG cluster (**Figure 5A right**).

**Figure 5.**
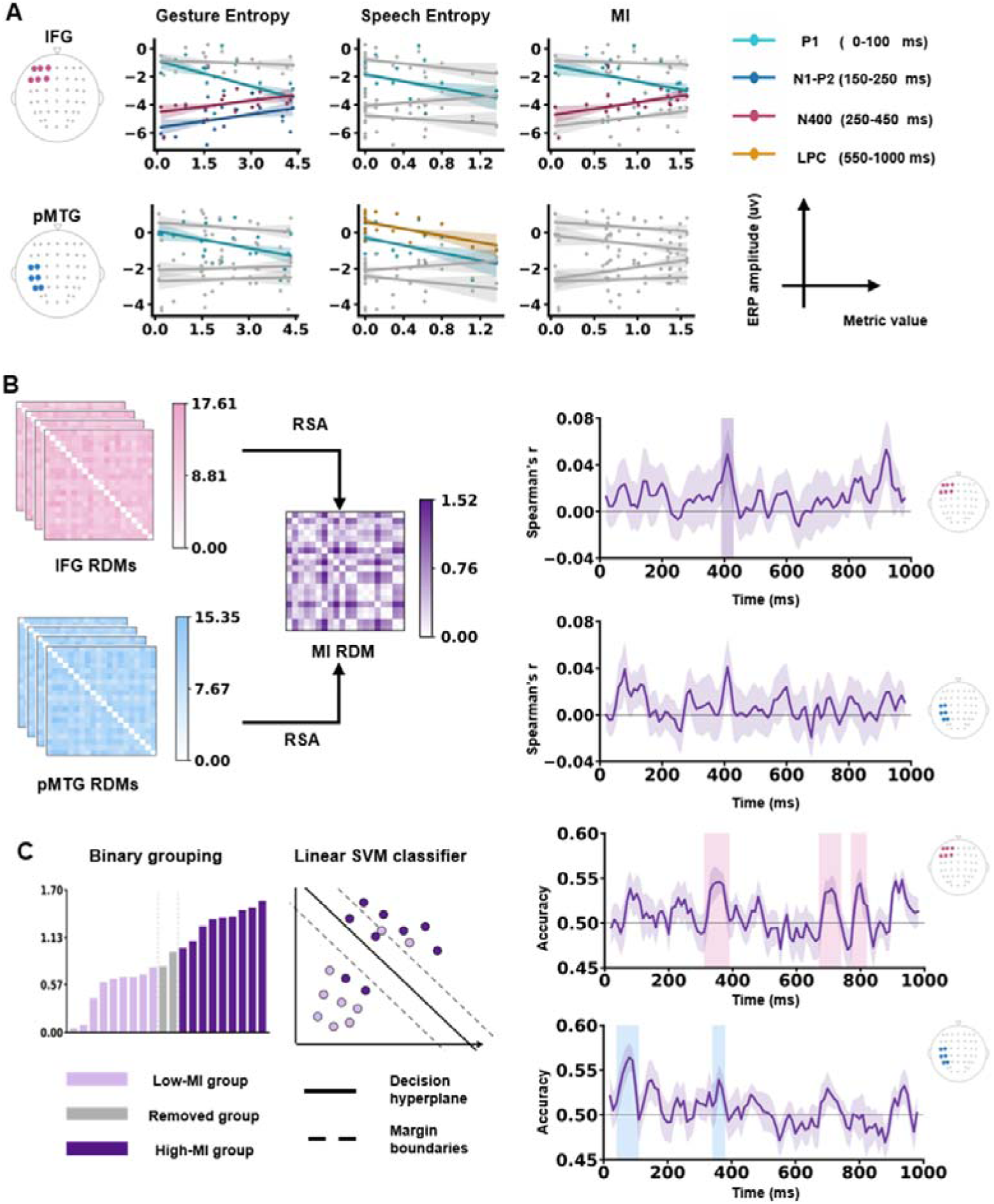
Temporal dynamics of information-theoretic representations during gesture– speech integration. **(A)** Linear mixed-effects model estimates of the associations between information-theoretic metrics and ERP amplitudes within predefined IFG- and pMTG-approximating electrode clusters. Gesture entropy, speech entropy, and MI were entered as stimulus-level predictors across P1, N1–P2, N400, and late positive component (LPC) time windows. Lines indicate model-estimated effects; gray lines indicate non-significant associations. **(B)** Time-resolved RSA of EEG patterns within IFG- and pMTG-approximating electrode clusters. Neural RDMs were constructed from multichannel EEG activity within sliding time windows and compared with the MI model RDM. Significant MI-related representational correspondence was identified within the IFG cluster during the N400 time window (390-430ms), whereas no significant effect was observed in the pMTG cluster. Shaded areas indicate significant temporal clusters identified using cluster-based permutation tests. **(C)** Time-resolved multivariate decoding of MI-related neural patterns. Stimuli were divided into low- and high-MI groups, and linear support vector machine (SVM) classifiers were trained to discriminate MI categories based on EEG activity patterns. Significant above-chance decoding was observed within both IFG- and pMTG-approximating electrode clusters across distinct temporal windows. Shaded regions indicate significant decoding clusters identified using cluster-based permutation tests.

Gesture entropy showed a broader modulation pattern within IFG. Higher gesture entropy was associated with reduced P1 amplitude (β = −0.783, SE = 0.172, *t* = −4.560, *p* < 0.001), followed by enhanced positivity during both the N1/P2 (β = 0.405, SE = 0.172, *t* = 2.348, *p* = 0.038) and the N400 windows (β = 0.360, SE = 0.163, *t* = 2.200, *p* = 0.038). In pMTG, gesture entropy selectively modulated early activity, producing a negative association with P1 amplitude (β = −0.410, SE = 0.146, *t* = −2.811, *p* = 0.021), with no reliable modulation at later stages (**Figure 5A left**).

Speech entropy demonstrated a distinct posterior temporal profile. In IFG, speech entropy was associated only with reduced P1 amplitude (β = −0.493, SE = 0.174, *t* = −2.830, *p* = 0.020). In pMTG, however, speech entropy influenced both early and late processing stages, producing reduced P1 (β = −0.426, SE = 0.146, *t* = −2.922, *p* = 0.007) and LPC amplitude (β = −0.384, SE = 0.125, *t* = −3.079, *p* = 0.007) (**Figure 5A middle**).

Together, these results reveal partially dissociable temporal profiles for modality-specific entropy and cross-modal MI. Early ERP modulation was observed across multiple information dimensions, suggesting sensitivity to stimulus-level information variability during initial processing stages. In contrast, later effects differentiated MI-related and entropy-related processing, with MI selectively modulating IFG activity during the semantic integration period and speech entropy showing a stronger influence on late pMTG responses.

Whole-scalp analyses revealed comparable temporal patterns for MI- and entropy-related effects, further supporting the robustness of these information-theoretic dynamics (Supplementary figure 2).

#### MI shapes IFG representational geometry during the N400-associated integration window

ERP analyses characterize changes in response magnitude but do not determine whether the relationships among stimuli are organized according to information-theoretic structure. We therefore applied time-resolved RSA to examine whether neural representational geometry preserved the stimulus relationships predicted by information-theoretic models. Within the predefined IFG and pMTG electrode clusters, neural RDMs were calculated across sliding time windows and compared with model RDMs derived from gesture entropy, speech entropy, and MI.

In the IFG electrode region, representational geometry significantly corresponded to the MI structure during the N400 time window (390–430 ms, mean *r* = 0.0495, mean *t* = 2.249, cluster-level *p* = 0.031) (**Figure 5B right top**). No significant MI-related RSA clusters were identified within the pMTG cluster (**Figure 5B right bottom**). Similarly, gesture entropy and speech entropy did not show significant ROI-level RSA effects (**Supplementary figure 3 top**). The timing of the IFG MI-related RSA effect overlapped with the MI-related ERP modulation observed in amplitude analyses and corresponded to the period during which fMRI identified MI-sensitive IFG representations. Whole-scalp RSA further revealed MI-related representational effects during early, and N400 time windows, with the N400 effect localized to frontal-central electrodes overlapping with the predefined IFG cluster (**Supplementary figure 1B left**). Thus, EEG RSA provides temporal evidence that IFG representations preserve the relational structure associated with gesture–speech MI during semantic integration.

#### Distributed temporal encoding of MI across IFG and pMTG

To complement RSA, which assesses whether neural representations preserve stimulus relationships, we performed time-resolved multivariate decoding to determine whether neural patterns contained sufficient information to distinguish stimuli with relatively high versus low values of each information-theoretic dimension.

In the IFG cluster, MI-related decoding emerged across multiple temporal windows. Significant above-chance decoding was observed between 310–390 ms (mean accuracy = 0.543, mean *t* = 2.784, cluster-level *p* = 0.006), 670–740 ms (mean accuracy = 0.535, *t* = 2.367, *p* = 0.015), and 770–820 ms (mean accuracy = 0.542, *t* = 2.537, *p* = 0.021) (**Figure 5C right top**). The earliest IFG decoding window (310–390 ms) overlapped with both the N400-related MI modulation and the RSA-sensitive period, indicating that IFG encodes MI both as changes in response magnitude and as distributed multichannel patterns. Later decoding clusters suggest that MI-related representations persist beyond the initial integration period.

In the pMTG cluster, MI decoding was detected during an early window (40–110 ms; mean accuracy = 0.557, *t* = 3.509, *p* = 0.001) and a later integration-related window (340–380 ms; mean accuracy = 0.539, *t* = 2.443, *p* = 0.024) (**Figure 5B right bottom)**. The later pMTG decoding window overlapped temporally with MI-related effects identified in fMRI and ERP analyses, supporting a role for posterior temporal cortex in representing gesture-speech correspondence.

Together, these decoding results demonstrate that MI-related information is dynamically represented across frontal and temporal regions. The temporal profiles observed in EEG complement the spatial organization identified by fMRI: whereas fMRI revealed MI-related representations within IFG and pMTG, EEG further revealed when these representations emerge and how their dynamics differ across regions. Specifically, IFG showed sustained MI-related encoding extending through the semantic integration period and later processing stages, whereas pMTG exhibited temporally distinct early and integration-related decoding windows.

Supplementary decoding analyses further examined whether gesture and speech entropy were represented at the neural level. These analyses revealed entropy-related decoding effects during selected temporal windows (**Supplementary figure 4**), suggesting that modality-specific information also influences transient neural dynamics during gesture– speech processing.

### Summary

Together, these EEG findings reveal distinct temporal profiles for modality-specific entropy and cross-modal MI during gesture–speech semantic integration. Entropy measures showed temporally specific effects associated with modality-related processing, with gesture and speech entropy exhibiting partially dissociable temporal signatures across frontal and temporal regions. In contrast, MI-related representations became prominent during the N400-associated integration period, with IFG exhibiting MI-consistent representational geometry and both IFG and pMTG showing discriminable MI-related neural patterns.

Source localization analyses further suggested that N400 MI-related activity involved frontal and temporal regions overlapping with the IFG–pMTG network identified in the fMRI analyses (**Supplementary figure 5**). Together with the fMRI findings, these results demonstrate complementary temporal and spatial signatures of information-theoretic representations: EEG captures the dynamic emergence of neural representations during stimulus processing, whereas fMRI identifies their spatial organization within the semantic network. Notably, MI showed consistent representational signatures across both temporal and spatial measurements, whereas entropy-related effects were primarily reflected in temporally specific EEG dynamics.

Together, these findings provide temporal evidence that cross-modal information structure emerges during gesture–speech integration and complement the spatially resolved fMRI evidence that MI organizes representations within the IFG–pMTG network.

### Combined analysis of Experiments 2 and 3

#### EEG–fMRI representational alignment reveals complementary temporal and spatial signatures of MI

The fMRI and EEG experiments provided complementary perspectives on MI-related representations during gesture–speech semantic integration. fMRI identified where MI-related representations were organized within the IFG–pMTG network, whereas EEG revealed when these representations emerged during semantic processing. However, whether these spatial and temporal signatures reflect a shared neural representation or distinct aspects of the same information-theoretic structure remains unclear.

To address this question, we performed cross-modal representational alignment analyses between EEG and fMRI. The EEG representation was derived from the frontal-central 390– 430 ms time window identified by EEG RSA, during which neural representational geometry significantly corresponded to the MI model. The fMRI representation was extracted from the MI-sensitive IFG searchlight cluster identified by fMRI RSA. This approach allowed us to examine whether MI-related representations identified across temporal and spatial measurements reflected overlapping or complementary representational structures.

#### EEG and fMRI independently capture MI-related representational structure

We first examined whether EEG and fMRI representations independently reflected the stimulus-level structure predicted by MI.

The EEG RDM was significantly correlated with the MI model RDM (Spearman *r* = 0.207, permutation *p* = 0.022), indicating that the temporal neural patterns captured during the N400 window preserved the stimulus-level structure predicted by MI. Similarly, the fMRI RDM extracted from the MI-sensitive IFG cluster was significantly correlated with the MI model RDM (*r* = 0.184, *p* = 0.035), demonstrating that local multivoxel activity patterns within IFG also reflected the same MI organization.

However, the EEG and fMRI RDMs were not significantly correlated with each other (*r* = −0.019, *p* = 0.865). Thus, although both modalities captured MI-related representational structure, they did not exhibit identical representational geometries (**Figure 6A)**.

**Figure 6.**
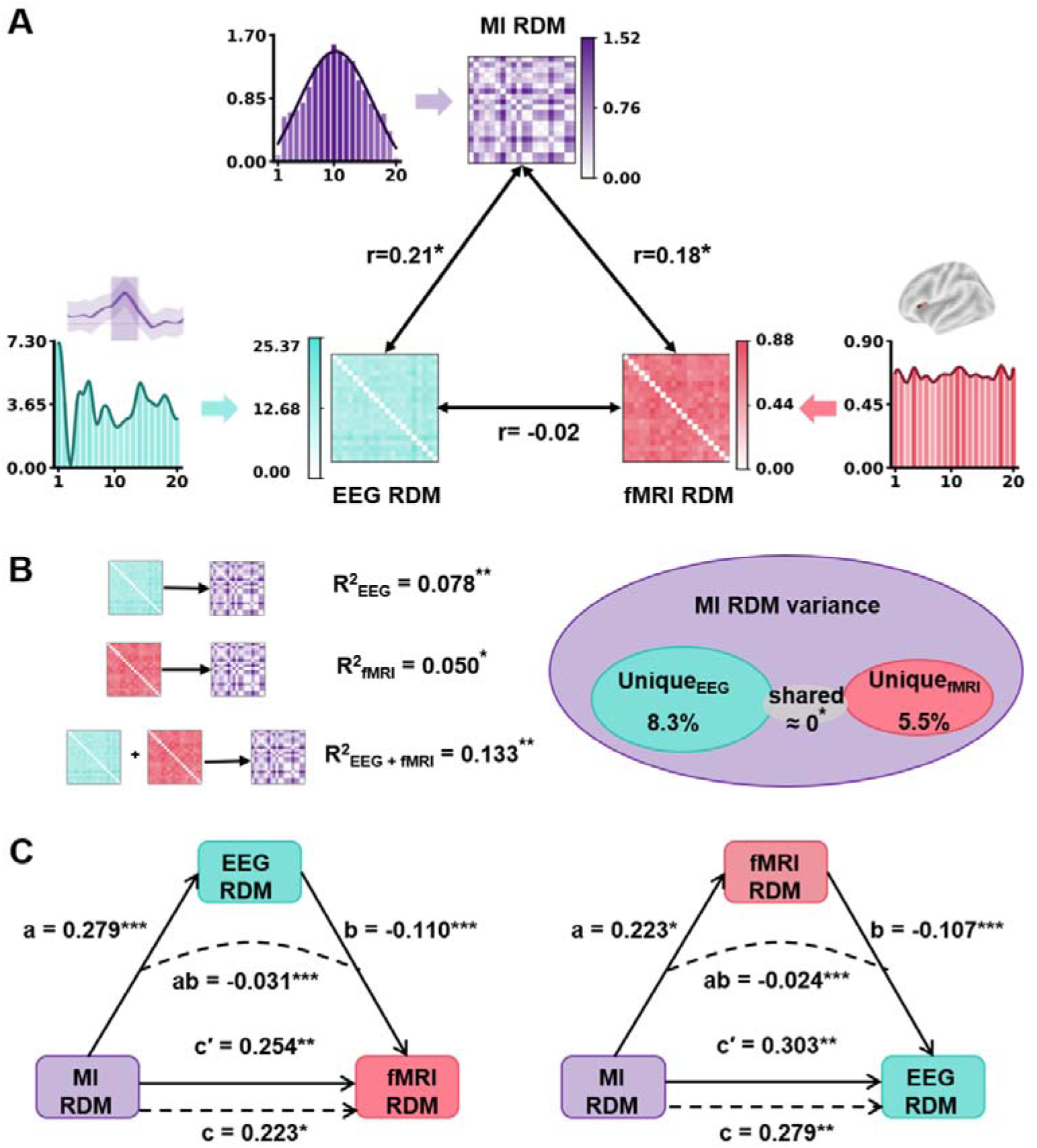
Cross-modal representational analyses reveal complementary EEG and fMRI signatures of MI. **(A)** Cross-modal representational alignment among EEG, fMRI, and the MI model. Group-level EEG and fMRI RDMs were extracted from the significant MI-related EEG RSA cluster (390-430 ms) and the MI-sensitive IFG fMRI cluster, respectively. Pairwise correlations were calculated among EEG, fMRI, and MI model RDMs to assess shared representational structure across measurement modalities. **(B)** Variance decomposition of MI-related representational structure. EEG and fMRI RDMs were used to predict variance in the MI model RDM. Variance explained by EEG, fMRI, and their combined representation (**Left**). Partitioning of explained variance into unique EEG, unique fMRI, and shared contributions (**Right**). **(C)** Representational decomposition was performed within representational similarity space to characterize the relationships among MI model, EEG, and fMRI representations. EEG representation was tested as a statistical mediator of the relationship between MI and fMRI representations (**Left**). fMRI representation was tested as a statistical mediator of the relationship between MI and EEG representations (**Right**). Standardized path coefficients are shown. * *p* < 0.05, ** *p* < 0.01, ** *p* < 0.001 after FDR correction.

These findings indicate that EEG and fMRI converge on the same information-theoretic dimension while capturing distinct neural signatures. EEG reflects the temporal evolution of MI-related representations during semantic integration, whereas fMRI reveals their spatial organization within the IFG network.

#### EEG and fMRI provide unique contributions to MI-related representations

To quantify whether EEG and fMRI captured overlapping or independent components of MI-related representational structure, we performed variance decomposition analysis using the MI model RDM as the target representation.

Both EEG and fMRI representations independently explained significant variance in the MI model. The EEG representation accounted for significant variance (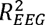 = 0.078, *p* = 0.007), and the fMRI representation also explained significant variance (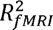 = 0.050, *p* = 0.018). When both representations were considered together, they explained additional variance beyond either modality alone (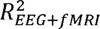 = 0.133, *p* = 0.001) (**Figure 6B left)**.

Variance partitioning further revealed significant unique contributions from both modalities. The unique contribution of EEG was significant (*Unique_EEG_* = 0.083, *p* = 0.005), as was the unique contribution of fMRI (*Unique_FMRI_* = 0.055, *p* = 0.012). In contrast, shared variance between EEG and fMRI representations was minimal (*shared*= −0.005, *p* = 0.045) (**Figure 6B right)**.

Together with the absence of significant EEG–fMRI RDM correlation, these results indicate that EEG and fMRI provide complementary information about MI-related representational organization. EEG captures temporally evolving aspects of MI representation, whereas fMRI captures spatially localized aspects within the semantic integration network.

#### Representational decomposition reveals distinct residual signatures of EEG and fMRI representations

Although EEG and fMRI independently reflected MI-related structure, their lack of direct representational correspondence raised the question of whether each modality contained distinct residual information beyond the shared MI component. To address this issue, we performed representational decomposition analyses within the RDM space.

When MI was specified as the predictor representation, EEG as the mediator, and fMRI as the outcome representation, MI was significantly associated with both EEG (path a = 0.279, *p* = 0.010) and fMRI representations (total effect c = 0.223, *p* = 0.020). The direct association between MI and fMRI remained significant after controlling for EEG (c′ = 0.254, *p* = 0.006). In contrast, the residual association between EEG and fMRI after controlling for MI was negative (path b = −0.110, *p* < 0.001), yielding a small but significant negative indirect effect (ab = −0.031, *p* < 0.001) (**Figure 6C left)**.

The complementary model, with fMRI specified as the mediator and EEG as the outcome representation, showed a similar pattern. MI was significantly associated with fMRI (path a = 0.223, *p* = 0.020), and EEG representations (total effect c = 0.279, *p* = 0.010). The direct association between MI and EEG remained significant after controlling fMRI (c′ = 0.303, *p* = 0.006) whereas the residual association between fMRI and EEG after accounting for MI was again negative (path b = −0.107, *p* < 0.001). This resulted in a small but significant negative indirect effect (ab = −0.024, *p* < 0.001) (**Figure 6C right)**.

The negative indirect effects should not be interpreted as inhibitory neural interactions. Instead, they indicate that after removing the variance explained by the MI model, EEG and fMRI contain statistically distinct residual representational structures. This interpretation is consistent with the variance decomposition results showing significant unique contributions and minimal shared variance.

### Summary

Cross-modal representational analyses demonstrated that EEG and fMRI converge on MI as a common information-theoretic dimension while capturing distinct aspects of its neural organization. Both EEG and fMRI representations significantly reflected the stimulus-level structure predicted by MI, indicating that MI-related stimulus structure is preserved across temporal and spatial measurements. However, the absence of significant EEG – fMRI representational correspondence and the variance decomposition results showing significant modality-specific contributions indicate that the two modalities do not reflect an identical neural representation.

Instead, EEG and fMRI provide complementary perspectives on MI-related organization. EEG captures the temporal emergence and evolution of MI-related representations during semantic integration, whereas fMRI identifies their spatial organization within the IFG–pMTG semantic network. Together, these findings indicate that MI is represented across neural timescales through distinct but convergent representational signatures rather than a shared neural code.

## Discussion

Through mathematical quantification of gesture and speech information using entropy and mutual information (MI), we investigated how information-theoretic structure shapes the neural organization of multisensory semantic integration across complementary experimental approaches. Causal perturbation using tDCS demonstrated that modulating activity across the left IFG and pMTG altered behavioral sensitivity to gesture–speech MI, establishing the functional relevance of this network for MI-related integration processes (**Experiment 1**).

Extending these causal findings, fMRI revealed the spatial organization of MI-related representations across the IFG and pMTG (**Experiment 2**), whereas EEG characterized the temporal dynamics of both modality-specific entropy and MI-related representations during semantic processing (**Experiment 3**). Finally, cross-modal representational analyses showed that EEG and fMRI converged on MI as a shared information-theoretic dimension while capturing complementary temporal and spatial signatures. Together, these findings reveal a progressive organization of multisensory semantic integration, in which neural responses and representational structures systematically track variation in stimulus-level information-theoretic metrics, with modality-specific entropy expressed through transient dynamics and MI emerging as a more consistent signature of cross-modal integration across neural scales.

It is widely acknowledged that a single, amodal system mediates the interactions among perceptual representations of different modalities^5,32,33^. Moreover, observations have suggested that semantic dementia patients experience increasing overregularization of their conceptual knowledge due to the progressive deterioration of this amodal system^34^. Consistent with this, the present study provides robust evidence, through the application of HD-tDCS, that the integration hubs for gesture and speech—the pMTG and IFG—operate in an incremental manner. This is supported by the progressive inhibition effect observed in these brain areas as MI of gesture and speech advances, indicating their functional contribution to cross-modal semantic processing.

The fMRI findings further clarify how MI-related representations are spatially organized across the IFG and pMTG. Although both IFG and pMTG contained MI-related information, they expressed this information through partially distinct representational properties. The IFG preserved the representational structure captured by MI, indicating that IFG activity patterns retain stimulus-level relationships defined by shared gesture–speech information. This finding complements previous accounts of IFG as a key region involved in semantic control and selection^35,36^, by suggesting that IFG also contributes to representing structured relationships among distributed semantic inputs. In contrast, pMTG patterns showed stronger discrimination of MI-defined stimulus differences, consistent with its proposed role as a convergence region linking perceptual and conceptual representations^37^.

Importantly, representational connectivity analyses demonstrated that IFG and pMTG exhibited significant stimulus-level representational correspondence, consistent with their close anatomical and functional relationship within the semantic network^22,23^. However, the strength of this correspondence was not enhanced for high-MI stimuli and was not predicted by continuous variation in MI. These findings indicate that, although IFG and pMTG are coordinated during multimodal semantic processing, their contributions to MI-related representation are not determined by MI-dependent modulation of inter-regional coupling. Instead, IFG and pMTG appear to contribute complementary aspects of MI-related representation within a shared semantic network, with this functional differentiation emerging independently of changes in representational connectivity.

Furthermore, we disentangled the sub-processes of integration with high-temporal ERPs, when representations of gesture and speech were variously presented. Early P1-N1 and P2 sensory effects linked to perception and attentional processes^28,38^ was comprehended as a reflection of the early audiovisual gesture-speech integration in the sensory-perceptual processing chain^39^. Note that a semantic priming paradigm was adopted here to create a top-down prediction of gesture over speech. The observed negative correlation of the P1 effect with speech entropy and the negative correlation of the P1 and N1-P2 effect with gesture entropy suggest that the early interaction of gesture-speech information was modulated by both top-down gesture prediction and bottom-up speech processing. Additionally, the lexico-semantic effect of the N400 and the LPC were differentially mediated by top-down gesture prediction, bottom-up speech encoding and their interaction: the N400 was correlated with both the gesture entropy and MI, but the LPC was negatively correlated only with the speech entropy.

The varying contributions of unisensory gesture-speech information and the convergence of multisensory inputs, as reflected in the correlation between distinct ERP components, are consistent with recent models suggesting that multisensory processing involves parallel detection of modality-specific information and hierarchical integration across multiple neural levels^4,40^. These processes are further characterized by coordination across multiple temporal scales^41^. Building on this, the present study offers additional evidence that the multi-level nature of gesture-speech processing is statistically structured, as measured by information matrix of unisensory entropy and multisensory convergence index of MI, the input of either source would activate a distributed representation, resulting in progressively functioning neural responses.

Considering the temporal profiles of the ERP components within the predefined IFG- and pMTG-approximating electrode clusters, we further examined how information-theoretic representations unfold over time within regions implicated in gesture–speech semantic integration. These findings reveal temporally differentiated contributions of IFG and pMTG to different aspects of information processing. First, early sensory-perceptual processing indexed by the P1 component reflected sensitivity to modality-specific information variability^27,28^. Both gesture entropy and speech entropy were negatively associated with P1 responses across IFG and pMTG regions, suggesting that increased uncertainty within either modality influenced early neural processing of gesture–speech stimuli. Second, the gesture representation was activated in the pMTG during the N1-P2 window and further exerted a top-down modulation over the phonological processing of speech^42^. The higher the certainty of gesture is, a larger modulation of gesture would be made upon speech, as indexed by a smaller gesture entropy with an enhanced N1-P2 amplitude. Third, during the N400-associated semantic integration period^43^, both gesture entropy and MI modulated IFG N400 responses, with higher values associated with reduced N400-related activity. The reduced N400 amplitude with the increase of gesture entropy may reflect weaker semantic constraint from gesture when the representation of gesture was wildly distributed. In contrast, larger MI indicates larger overlapped neural populations activated by gesture and speech, indexing easier integration and less semantic conflict were observed, as indexed by reduced N400 responses. Last, the activated speech representation would disambiguate and reanalyze the semantic information and further unify into a coherent comprehension in the pMTG^12,44^. As speech entropy increases, indicating greater uncertainty in the information provided by speech, more cognitive effort is directed towards selecting the targeted semantic representation. This leads to enhanced involvement of the pMTG and a corresponding reduction in LPC amplitude.

It should be noted that the association between ERP components and cortical regions described above was assessed by examining temporally specific EEG responses within IFG-and pMTG-approximating electrode clusters. Neither gesture nor speech entropy showed reliable spatial representations in fMRI. This difference likely reflects the differential sensitivity of EEG and fMRI to entropy- and MI-related representations, highlighting distinct temporal and spatial properties of information processing during multisensory integration. Specifically, modality-specific information dynamics may be more transient and less stable than the cross-modal representational structures captured by MI. Importantly, EEG-based regional associations should be interpreted as temporal dynamics expressed within electrode clusters approximating IFG and pMTG rather than as direct evidence of cortical localization. Thus, fMRI identifies where consistent MI-related representations are organized within the semantic network, whereas EEG reveals when modality-specific and cross-modal information dynamics emerge during processing.

MI showed consistent representations across both EEG and fMRI, supporting its role as a more stable cross-modal information structure. Nevertheless, the representational structures identified by the two modalities showed limited direct correspondence, indicating that EEG and fMRI do not reflect an identical neural code. Instead, these findings suggest that the two approaches capture complementary dimensions of the same information-processing framework. EEG primarily reveals the temporal evolution of information representations, including the emergence of MI-related structure during the N400-associated integration period, whereas fMRI captures the spatial organization of these representations within the IFG–pMTG semantic network. Therefore, MI should not be considered a fixed neural pattern localized to a specific region or processing stage, but rather a computational structure dynamically expressed across different neural scales. The convergence between EEG and fMRI supports a cross-scale framework in which multisensory semantic integration is progressively organized through coordinated temporal dynamics and spatially distributed representations.

Note that the cortical involvement and ERP components discussed above are derived from a deliberate alignment of speech onset with gesture DP, creating an artificial priming effect with gesture semantically preceding speech. Caution is advised when generalizing these findings to the spontaneous gesture-speech relationships, although gestures naturally precede speech^45^. The information-theoretic framework used here extends traditional MI^46^ by representing gesture and speech in a shared probabilistic semantic space with reliability-weighted alignment, allowing graded quantification of cross-modal semantic relationships beyond simple label overlap. However, because the current MI formulation quantifies shared information between aligned semantic dimensions, future extensions to complementary or non-overlapping gesture–speech relationships^47^ may require additional metrics that capture semantic complementarity beyond shared representations.

Limitations exist. ERP components and cortical engagements were linked through intermediary variables of entropy and MI. Dissociations were observed between ERP components and cortical engagement. Importantly, there is no direct evidence of the brain structures underpinning the corresponding ERPs, necessitating clarification in future studies. Additionally, the current study incorporated a restricted set of entropy and MI measures. The generalizability of the findings should be assessed in future studies using a more extensive range of matrices.

In summary, this study establishes an information-theoretic framework for investigating multisensory semantic integration, showing that semantic representations are shaped by the statistical structure of distributed sensory information. This framework provides a principled approach for linking computational properties of stimuli with their neural implementation and offers broader insights into how the brain constructs coherent meaning from heterogeneous inputs.

## Material and methods

### Participants

One hundred and two young Chinese participants signed written informed consent forms and took part in the present study (Experiment 1: 29 females, 23 males, age = 20 ± 3.40 years; Experiment 2: 15 females, 13 males, age = 26 ± 2.95 years; Experiment 3: 12 females, 10 males, age = 21 ± 3.53 years). All of the participants were right-handed, had normal or corrected-to-normal vision and were paid ¥100 per hour for their participation. All experiments were approved by the Ethics Committee of the Institute of Psychology, Chinese Academy of Sciences.

### Stimuli

Twenty gestures (**Supplementary Table 1**) with 20 semantically congruent speech signals taken from previous study^23^ were used. The stimuli set were recorded from two native Chinese speakers (1 male, 1 female). To validate the stimuli, 30 participants were recruited to replicate the multisensory index of semantic congruency effect, hypothesizing that reaction times for semantically incongruent gesture-speech pairs would be significantly longer than those for congruent pairs. The results confirmed this hypothesis, with a significantly (*t*(29) = 7.16, *p* < 0.001) larger reaction time when participants were asked to judge the gender of the speaker if gesture contained incongruent semantic information with speech (a ‘cut’ gesture paired with speech word ‘喷 pen1 (spray)’: mean = 554.51 ms, SE = 11.65) relative to when they were semantically congruent (a ‘cut’ gesture paired with ‘剪 jian3 (cut)’ word: mean = 533.90 ms, SE = 12.02)^23^.

Additionally, two separate pre-tests with 30 subjects in each (pre-test 1: 16 females, 14 males, age = 24 ± 4.37 years; pre-test 2: 15 females, 15 males, age = 22 ± 3.26 years) were conducted to determine the comprehensive values of gesture and speech. Participants were presented with segments of increasing duration, beginning at 40 ms, and were prompted to provide a single verb to describe either the isolated gesture they observed (pre-test 1) or the isolated speech they heard (pre-test 2). For each gesture or speech, the action verb consistently provided by participants across four to six consecutive repetitions—with the number of repetitions varied to mitigate learning effects—was considered the comprehensive response for the gesture or speech. The initial instance duration was marked as the discrimination point (DP) for gesture (mean = 183.78 ± 84.82ms) or the identification point (IP) for speech (mean = 176.40 ± 66.21ms) (**Figure 1A top**).

To quantify information content, comprehensive responses for each item were converted into Shannon’s entropy (H) as a measure of information richness (**Figure 1A bottom**). With no significant gender differences observed in both gesture (*t*(20) = 0.21, *p* = 0.84) and speech (*t*(20) = 0.52, *p* = 0.61), responses were aggregated across genders, resulting in 60 answers per item (**Supplementary Table 2**). Here, p(xi) and p(yi) represent the distribution of 60 answers for a given gesture (**Supplementary Table 2B**) and speech (**Supplementary Table 2A**), respectively. High entropy indicates diverse answers, reflecting broad representation, while low entropy suggests focused lexical recognition for a specific item (**Figure 2B**). MI was used to measure the overlap between gesture and speech information, calculated by subtracting the entropy of the combined gesture-speech dataset (Entropy(gesture + speech)) from the sum of their individual entropies (Entropy(gesture) + Entropy(speech)) (see **Supplementary Table 2C**). For specific gesture-speech combinations, equivalence between the combined entropy and the sum of individual entropies (gesture or speech) indicates absence of overlap in response sets. Conversely, significant overlap, denoted by a considerable number of shared responses between gesture and speech datasets, leads to a noticeable discrepancy between combined entropy and the sum of gesture and speech entropies. Elevated MI values thus signify substantial overlap, indicative of a robust mutual interaction between gesture and speech.

Additionally, the number of responses provided for each gesture and speech, as well as the total number of combined responses, were also recorded. The quantitative data for each stimulus, including gesture entropy, speech entropy, joint entropy, MI, and the respective counts, are presented in **Supplementary Table 3.**

To determine whether entropy or MI values corresponds to distinct neural changes, the current study first aggregated neural responses (including inhibition effects of tDCS and TMS or ERP amplitudes) that shared identical entropy or MI values, prior to conducting correlational analyses.

### Experimental procedure

Given that gestures induce a semantic priming effect on concurrent speech^48^, this study utilized a semantic priming paradigm in which speech onset was aligned with the DP of each gesture^23,48^, the point at which the gesture transitions into a lexical form^45^. The gesture itself began at the stroke phase, a critical moment when the gesture conveys its primary semantic content.

An irrelevant factor of gender congruency (e.g., a man making a gesture combined with a female voice) was created^22,23,49^. This involved aligning the gender of the voice with the corresponding gender of the gesture in either a congruent (e.g., male voice paired with a male gesture) or incongruent (e.g., male voice paired with a female gesture) manner. This approach served as a direct control mechanism, facilitating the investigation of the automatic and implicit semantic interplay between gesture and speech. In light of previous findings indicating a distinct TMS-disruption effect on the semantic congruency of gesture-speech interactions^23^, both semantically congruent and incongruent pairs were included in Experiment 1. Experiment 2 and Experiment 3, conversely, exclusively utilized semantically congruent pairs to elucidate ERP metrics indicative of nuanced semantic progression.

Gesture–speech pairs were presented randomly using Presentation software (www.neurobs.com). Participants were asked to look at the screen but respond with both hands as quickly and accurately as possible merely to the gender of the voice they heard. The RT and the button being pressed were recorded. The experiment started with a fixation cross presented on the center of the screen, which lasted for 0.5-1.5 sec.

### Experiment 1: HD-tDCS protocol and data analysis

Participants were divided into two groups, with each group undergoing HD-tDCS stimulation at different target sites (IFG or pMTG). Each participant completed three experimental sessions, spaced one week apart, during which 480 gesture-speech pairs were presented across various conditions. In each session, participants received one of three types of HD-tDCS stimulation: Anodal, Cathodal, or Sham. The order of stimulation site and type was counterbalanced using a Latin square design to control for potential order effects.

HD-tDCS protocol employed a constant current stimulator (The Starstim 8 system) delivering stimulation at an intensity of 2mA. A 4 * 1 ring-based electrode montage was utilized, comprising a central electrode (stimulation) positioned directly over the target cortical area and four return electrodes encircling it to provide focused stimulation. Building on a meta-analysis of prior fMRI studies examining gesture-speech integration^22^, we targeted Montreal Neurological Institute (MNI) coordinates for the left IFG at (−62, 16, 22) and the pMTG at (−50, −56, 10). In the stimulation protocol for HD-tDCS, the IFG was targeted using electrode F7 as the optimal cortical projection site^50^, with four return electrodes placed at AF7, FC5, F9, and FT9. For the pMTG, TP7 was selected as the cortical projection site^50^, with return electrodes positioned at C5, P5, T9, and P9. The stimulation parameters included a 20-minute duration with a 5-second fade-in and fade-out for both Anodal and Cathodal conditions. The Sham condition involved a 5-second fade-in followed by only 30 seconds of stimulation, then 19’20 minutes of no stimulation, and finally a 5-second fade-out (**Figure 1B**). Stimulation was controlled using NIC software, with participants blinded to the stimulation conditions.

All incorrect responses (702 out of the total number of 24960, 2.81% of trials) were excluded. To eliminate the influence of outliers, a 2SD trimmed mean for every participant in each session was also calculated. To examine the relationship between the degree of information and neural responses, we conducted Pearson correlation analyses using a sample of 20 sets. Neural responses were quantified based on the effects of HD-tDCS (active tDCS minus sham tDCS) on the semantic congruency effect, defined as the difference in reaction times between semantic incongruent and congruent conditions (Rt(incongruent) - Rt(congruent)). This effect served as an index of multisensory integration^49^ within the left IFG and pMTG. The variation in information was assessed using three information-theoretic metrics. To account for potential confounds related to multiple candidate representations, we conducted partial correlation analyses between the tDCS effects and gesture entropy, speech entropy, and MI, controlling for the number of responses provided for each gesture and speech, as well as the total number of combined responses. Given that HD-tDCS induces overall disruption at the targeted brain regions, we hypothesized that the neural activity within the left IFG and pMTG would be progressively affected by varying levels of multisensory convergence, as indexed by MI. Moreover, we hypothesized that the modulation of neural activity by MI would differ between the left IFG and pMTG, as reflected in the differential modulation of response numbers in the partial correlations, highlighting their distinct roles in semantic processing^44^. False discovery rate (FDR) correction was applied for multiple comparisons.

### Experiment 2: fMRI data acquisition, preprocessing and representational analyses

#### fMRI data acquisition

Functional and structural MRI data were acquired using a 3T Siemens Prisma MRI scanner equipped with a 64-channel head coil. High-resolution T1-weighted anatomical images were collected using a magnetization-prepared rapid acquisition gradient echo (MPRAGE) sequence (0.8 mm isotropic voxels; repetition time (TR) = 2300 ms; echo time (TE) = 2.07 ms; inversion time (TI) = 1100 ms; flip angle = 8°; in-plane matrix = 320 × 280; 240 slices; GRAPPA acceleration factor = 2).

Task-related blood-oxygen-level-dependent (BOLD) functional images were acquired using a T2*-weighted gradient-echo echo-planar imaging (EPI) sequence (TR = 1500 ms; TE = 30 ms; flip angle = 90°; slice thickness = 2 mm; in-plane matrix size = 96 × 96). Functional images were collected across four task runs, with each run lasting approximately 10 min.

#### fMRI data preprocessing

Functional image preprocessing was performed using SPM12 (Wellcome Trust Centre for Neuroimaging, London, UK). Functional images were first corrected for slice acquisition. Images were subsequently realigned and resliced to correct for head motion. Functional runs within each participant were jointly realigned using a six-parameter rigid-body transformation, with all volumes aligned to the participant-specific mean functional image. The six motion parameters were retained as nuisance regressors in subsequent first-level analyses.

The individual T1-weighted anatomical image was coregistered to the mean functional image using normalized mutual information. Spatial normalization was performed using SPM12’s unified segmentation-normalization procedure. Nonlinear deformation fields were estimated from anatomical tissue segmentation and subsequently applied to functional images. Functional images were resampled into Montreal Neurological Institute (MNI) standard space at an isotropic voxel size of 2 × 2 × 2 mm³ using fourth-order B-spline interpolation. Following spatial normalization, functional images were smoothed using a 6-mm full-width-at-half-maximum Gaussian kernel and subsequently used for all downstream analyses.

#### Trial-wise beta estimation and region-of-interest (ROI) definition

First-level general linear models (GLMs) were constructed separately for each participant using SPM12. Individual stimulus presentations were modeled as event-related regressors and convolved with the canonical hemodynamic response function. Motion parameters were included as nuisance regressors, and a high-pass filter with a 128s cutoff was applied to remove low-frequency signal drifts.

To estimate stimulus-specific neural representations, single-trial beta estimates were generated using a least-square-all (LSA) approach^51^. Each stimulus presentation was modeled as a separate regressor in a single GLM. The resulting stimulus-specific beta estimates were used for all multivariate analyses, including representational similarity analysis (RSA), multivoxel pattern analysis (MVPA), and representational connectivity analysis.

ROIs were defined independently from the representational analyses based on significant clusters identified from a whole-brain group-level contrast comparing semantically incongruent and congruent trials. This independent ROI definition was used to identify regions involved in multimodal semantic processing while avoiding circularity in subsequent representational analyses. Significant whole-brain clusters were transformed into ROI masks in MNI space and parcellated according to the Brainnetome Atlas. The left inferior frontal gyrus (IFG) ROI corresponded to BNA37 (A44op_L; pars opercularis), whereas the left posterior middle temporal gyrus (pMTG) ROI corresponded to BNA75 (A22c_L).

Because the aim of the representational analyses was to characterize information structures supporting multimodal semantic integration, only correct trials from the semantically congruent condition were included. Trials with reaction times exceeding ±3 standard deviations from each participant’s mean reaction time were excluded before analysis.

### Searchlight representational similarity analysis

Searchlight representational similarity analysis (RSA) was conducted within the independently defined left IFG and pMTG ROIs to determine whether local neural representational geometry reflected stimulus-level information-theoretic structure.

For each information-theoretic dimension (gesture entropy, speech entropy and MI), a model representational dissimilarity matrix (RDMs) was constructed based on pairwise Euclidean distances among stimulus values. Each model RDM represented the predicted relational structure among stimuli according to the corresponding information-theoretic measure.

Within each searchlight sphere, neural RDMs were generated from stimulus-specific beta estimates by calculating correlation distance between multivoxel activity patterns. Searchlight spheres were defined with a radius of three voxels. Correspondence between neural and model RDMs was quantified using Spearman rank correlations between the upper triangular elements of the two matrices.

Participant-level RSA maps were generated by assigning correlation coefficients to the center voxel of each searchlight sphere. Group-level statistical inference was performed separately for gesture entropy, speech entropy, and MI within each ROI using non-parametric permutation testing (5000 permutations) with cluster-level correction (*p* < 0.05). This approach allowed us to determine whether local neural representations preserved the relational structure associated with specific information-theoretic dimensions.

### ROI-based multivoxel pattern analysis

ROI-based multivoxel pattern analysis (MVPA) was performed within the left IFG and left pMTG ROIs to determine whether local neural activity patterns contained information that allowed discrimination between different levels of information-theoretic states.

For each information-theoretic dimension, stimuli were ranked according to their corresponding values. The nine stimuli with the lowest values were assigned to the low-information group, whereas the nine stimuli with the highest values were assigned to the high-information group. The two intermediate stimuli were excluded to maximize separation between classes while maintaining balanced classification.

This classification procedure was applied consistently across gesture entropy, speech entropy, and MI analyses. Because speech entropy showed a highly discrete distribution, with nine stimuli sharing identical entropy values of zero, these stimuli were assigned to the low-entropy group, whereas the nine stimuli with the highest entropy values were selected as the high-entropy group. MI values were available for 19 stimuli, with two stimuli showing identical MI values. MI-based analyses were therefore performed on these 19 stimuli using the same balanced high-versus-low classification strategy.

Stimulus-level labels were assigned to individual trial. Multivoxel features were extracted separately from each ROI and standardized within each cross-validation fold. Principal component analysis (PCA) was subsequently applied, retaining components explaining 99% of the variance, followed by linear support vector classification (LinearSVC; C = 1; class weight = balanced)^52^. All preprocessing steps were performed independently within each cross-validation fold to prevent information leakage.

Classification performance was evaluated using leave-one-run-out cross-validation. Balanced accuracy was used as the performance metric because of the binary classification design. At the group level, decoding accuracy was compared against chance level (0.5) using one-sample t-tests.

### IFG**–**pMTG representational connectivity analysis

To determine whether IFG and pMTG exhibited similar stimulus-level representational organization and whether such correspondence was modulated by information-theoretic properties, representational connectivity analyses were performed.

For each participant, stimulus-level neural RDMs were generated separately within IFG and pMTG ROIs using correlation distance. Representational connectivity was quantified as the Spearman correlation between the upper triangular elements of the two ROI-specific RDMs. Correlation coefficients were subsequently Fisher z-transformed before group-level statistical testing.

First, we tested whether IFG and pMTG showed significant representational correspondence across all stimuli. This analysis assessed whether the two regions organized stimulus representations in a similar manner during multimodal semantic processing.

Second, we examined whether representational connectivity was modulated by information-theoretic properties. Stimuli were divided into high- and low-information subsets according to gesture entropy, speech entropy, or MI values using the same ranking procedure applied in MVPA. Representational connectivity was calculated separately for each subset, and differences between high-versus low-information conditions were assessed using paired comparisons.

Finally, continuous modulation analyses were performed to determine whether stimulus-level information-theoretic values predicted IFG–pMTG representational connectivity. For each stimulus, representational connectivity values were correlated with gesture entropy, speech entropy, and MI values across stimuli. Group-level significance was assessed using one-sample t-tests.

### Experiment 3: Electroencephalogram (EEG) recording and data analysis EEG data acquisition and preprocessing

Experiment 3, comprising a total of 1760 gesture-speech pairs, was completed in a single-day session. EEG were recorded from 48 Ag/AgCl electrodes mounted in a cap according to the 10-20 system^38^, amplified with a PORTI-32/MREFA amplifier (TMS International B.V., Enschede, NL) and digitized online at 500 Hz (bandpass, 0.01-70 Hz). EEGLAB, a MATLAB toolbox, was used to analyze the EEG data^53^. Vertical and horizontal eye movements were measured with 4 electrodes placed above the left eyebrow, below the left orbital ridge and at bilateral external canthus. All electrodes were referenced online to the left mastoid. Electrode impedance was maintained below 5 KΩ. The average of the left and right mastoids was used for re-referencing. A high-pass filter with a cutoff of 0.05 Hz and a low-pass filter with a cutoff of 30 Hz were applied. Semi-automated artifact removal, including independent component analysis (ICA) for identifying components of eye blinks and muscle activity, was performed (**Figure 1D**). Participants with rejected trials exceeding 30% of their total were excluded from further analysis.

All incorrect responses were excluded (147 out of 1760, 8.35% of trials). To eliminate the influence of outliers, a 2 SD trimmed mean was calculated for every participant in each condition. Data were epoched from the onset of speech and lasted for 1000 ms. To ensure a clean baseline with no stimulus presented, a 200 ms pre-stimulus baseline correction was applied before gesture onset.

### ROI definition and ERP amplitude analysis

To characterize the temporal dynamics through which information-theoretic properties modulated multimodal semantic processing, two electrode clusters were defined as a priori based on their anatomical correspondence with the fMRI-defined gesture-speech integration network. The frontal electrode cluster (F5, F3, F1, FC5, FC3, FC1) was used as an approximation of the left IFG, whereas the centro-parietal temporal cluster (C5, C3, CP5, CP3, P5, P3) was used as an approximation of left pMTG^54^.

For each participant and stimulus, EEG amplitudes were averaged across electrodes within each ROI. Mean amplitudes were extracted from four predefined temporal windows corresponding to major stages of multimodal processing: the P1 effect observed within the time window of 0-100 ms^27,28^, the N1-P2 effect observed between 150-250ms^27,28^, the N400 within the interval of 250-450ms^14,28,29^, and the LPC spanning from 550-1000ms^30,31^.

For each ROI × time-window combination, linear mixed-effects models were fitted separately for gesture entropy, speech entropy, and mutual information:

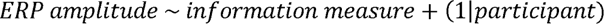

Each observation corresponded to the ERP response elicited by an individual stimulus within each participant. Information-theoretic variables were z-scored across the 20 stimuli before model fitting. The fixed-effect coefficient therefore represented the change in ERP amplitude associated with a one-standard-deviation increase in the corresponding information measure. Multiple comparisons were controlled using FDR correction across the four predefined time windows within each ROI × information-measure analysis.

These analyses were performed across all three information-theoretic dimensions to characterize both modality-specific uncertainty and cross-modal integration effects.

### Time-resolved representational similarity analysis

To determine whether neural activity preserved the continuous representational structure predicted by information-theoretic relationships, time-resolved RSA was performed within the predefined IFG and pMTG electrode clusters. MI was selected as the primary integration-related variable because it quantifies shared information between gesture and speech representations and directly captures cross-modal coupling strength. Complementary analyses using gesture entropy and speech entropy were conducted to characterize modality-specific uncertainty representations and are reported in Supplementary information.

For each participant and time window, stimulus-specific EEG patterns were extracted from the predefined electrode clusters. Neural RDMs were constructed using Euclidean distance between stimulus-level activity patterns.

The MI model RDM was constructed by calculating pairwise Euclidean distances between MI values across stimuli. Correspondence between neural and model RDMs was quantified using Spearman correlations between the upper triangular elements of the two matrices. Correlation coefficients were Fisher z-transformed and submitted to group-level one-sample t-tests against zero. Statistical significance was assessed using cluster-based permutation testing with sign-flipping correction.

### Time-resolved MI decoding

To determine when neural activity patterns contained information distinguishing different levels of cross-modal integration, time-resolved multivariate decoding was performed using MI as the classification variable.

For each participant, sliding-window EEG patterns were extracted from the predefined IFG and pMTG electrode clusters between 0 and 1000 ms after stimulus onset. A 40 ms sliding window was applied with a 10 ms step size. Stimuli were ranked according to MI values. The nine stimuli with the lowest MI values were assigned to the low-MI group, whereas the nine stimuli with the highest MI values were assigned to the high-MI group. The two intermediate stimuli were excluded to maximize class separation while maintaining balanced classification.

Single-trial EEG activity patterns within each ROI and time window were entered into a classification pipeline consisting of feature standardization, principal component analysis (PCA; retaining 99% explained variance), and linear support vector classifier (LinearSVC; C = 1, class weight = balanced). All preprocessing steps were performed independently within each cross-validation fold to avoid information leakage.

Classification performance was evaluated using leave-one-stimulus-out cross-validation. This procedure was selected to test whether classifiers generalized across individual stimulus identities rather than relying on repeated observations of the same stimulus. Balanced accuracy was used as the performance metric. At the group level, decoding accuracy at each time window was compared against chance level (0.5) using one-sample t-tests. Significant temporal clusters were identified using cluster-based permutation testing with sign-flipping across participants (*p* < 0.05).

### EEG–fMRI representational correspondence analysis

To determine whether EEG and fMRI captured convergent or complementary aspects of MI-related semantic representations, cross-modal representational analyses^55^ were performed between EEG and fMRI datasets.

Because EEG and fMRI data were acquired from independent participant samples, these analyses were conducted at the group-averaged representational level. Therefore, results were interpreted as comparisons of population-level representational geometry rather than subject-level neural coupling or directional interactions.

### Construction of EEG and fMRI representational matrices

For EEG data, representations were extracted from the frontal-central electrode cluster and temporal window showing significant MI-related RSA effects. EEG activity was averaged across selected electrodes and time points to generate a stimulus-level representation vector for each of stimuli. Pairwise Euclidean distances were calculated to construct a 20 × 20 EEG neural RDM.

For fMRI data, representations were extracted from the MI-sensitive searchlight cluster identified in the fMRI RSA analysis. Trial-wise beta estimates were averaged within this cluster for each stimulus, and pairwise correlation distances were calculated to construct the fMRI neural RDM. The MI model RDM was constructed using pairwise Euclidean distances between stimulus-level MI values.

### Cross-modal representational similarity analysis

To evaluate whether EEG and fMRI independently preserved MI-related stimulus structure, Spearman correlations were calculated between: (1) EEG neural RDM and MI model RDM; (2) fMRI neural RDM and MI model RDM; (3) EEG neural RDM and fMRI neural RDM. Statistical significance was assessed using permutation tests (5,000 iterations) by randomly shuffling stimulus labels while preserving the structure of the RDM.

### Variance decomposition analysis

To quantify the unique and shared contributions of EEG and fMRI representations to MI-related structure, representational variance decomposition was performed using the MI model RDM as the target representation.

Vectorized upper-triangular elements of EEG and fMRI RDMs were entered into regression models including: (1) EEG RDM alone; (2) fMRI RDM alone; (3) EEG and fMRI RDM jointly.

Explained variance (*R*^2^) of each model was calculated. Unique contributions were estimated as:

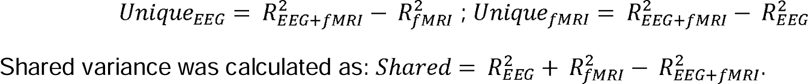

Statistical significance was assessed using permutation testing with stimulus-label randomization (5,000 iterations).

### Representational decomposition analysis

To further characterize the statistical relationship among MI, EEG and fMRI relationships, representational decomposition analyses were performed within the RDM vector space. These analyses were designed to determine whether EEG and fMRI captured overlapping or distinct components of MI-related representational structure. Because EEG and fMRI were obtained from independent participant samples, mediation effects were interpreted as statistical relationships among representational geometries rather than evidence for directional neural influence or temporal causality.

Two complementary models were tested: (1) MI model RDM→ EEG RDM → fMRI RDM; (2) MI model RDM→ fMRI RDM → EEG RDM. Analyses were conducted on the vectorized upper triangular elements of group-level RDMs. Standardized regression coefficients were estimated for four effects. The path a quantified the association between the MI model representation and the proposed mediator representation. Path b quantified the residual association between mediator and outcome representations after controlling MI. Path c represented the total effect of MI on the outcome representation, whereas the path c′ represented the direct effect after accounting for the mediator. The indirect effect was calculated as the product of coefficients (ab). Statistical significance of representational paths was assessed using permutation-based stimulus-label randomization (5,000 iterations).

## Acknowledgments

This research was supported by grants from the STI 2030—Major Projects 2021ZD0201500, the Beijing Natural Science Foundation (4262073), the National Natural Science Foundation of China (31822024, 31800964), the Scientific Foundation of Institute of Psychology, Chinese Academy of Sciences (E2CX3625CX), and the Strategic Priority Research Program of Chinese Academy of Sciences (XDB32010300).

## Author contributions

Conceptualization, W.Y.Z. and Y.D.; Investigation, W.Y.Z. and Z.Y.L.; Formal Analysis, W.Y.Z. and Z.Y.L.; Methodology, W.Y.Z. and Z.Y.L.; Validation, Z.Y.L. and X.L.; Visualization, W.Y.Z. and Z.Y.L. and X.L.; Funding Acquisition, W.Y.Z. and Y.D.; Supervision, Y.D.; Project administration, Y.D.; Writing – Original Draft, W.Y.Z.; Writing – Review & Editing, W.Y.Z., Z.Y.L., X.L., and Y.D.

## Data and Code Availability Statements

The data supporting the findings of this study are publicly available on the Open Science Framework (OSF) at https://doi.org/10.17605/OSF.IO/N82CG.

## SUPPLEMENTARY INFORMATION

**Supplementary Table 1.** Gesture description and paring with incongruent and congruent speech.

| **Gesture** | **Description** | **Congruent speech** | **Incongruent speech** |
| --- | --- | --- | --- |
| an4 (press) | press button | an4 (press) | yun4 (iron) |
| bai1 (break) | break chopsticks | bai1 (break) | yao2 (shake) |
| ca1 (wipe) | wipe desk | ca1 (wipe) | reng1 (throw) |
| ding1 (hammer) | hammer nail | ding1 (hammer) | tui1 (push) |
| feng2 (sew) | sew cloth | feng2 (sew) | ti2 (lift) |
| ji3 (squeeze) | squeeze sponge | ji3 (squeeze) | si1 (tear) |
| jian3 (cut) | cut paper | jian3 (cut) | sao1 (sweep) |
| jiao3 (stir) | stir flour | jiao3 (stir) | shan1 (slap) |
| ju4 (saw) | saw wood | ju4 (saw) | ning3 (twist) |
| ning3 (twist) | twist towel | ning3 (twist) | ju4 (saw) |
| pen1 (spray) | spray water | pen1 (spray) | qie1 (slice) |
| qie1 (slice) | slice fruit | qie1 (slice) | pen1 (spray) |
| reng1 (throw) | throw ball | reng1 (throw) | ca1 (wipe) |
| shan1 (slap) | slap face | shan1 (slap) | jiao3 (stir) |
| sao1 (sweep) | sweep floor | sao1 (sweep) | jian3 (cut) |
| si1 (tear) | tear paper | si1 (tear) | ji3 (squeeze) |
| ti2 (lift) | lift basket | ti2 (lift) | feng2 (sew) |
| tui1 (push) | push door | tui1 (push) | ding1 (hammer) |
| yao2 (shake) | shake bag | yao2 (shake) | bai1 (break) |
| yun4 (iron) | iron cloth | yun4 (iron) | an4 (press) |

**Supplementary Table 2.**
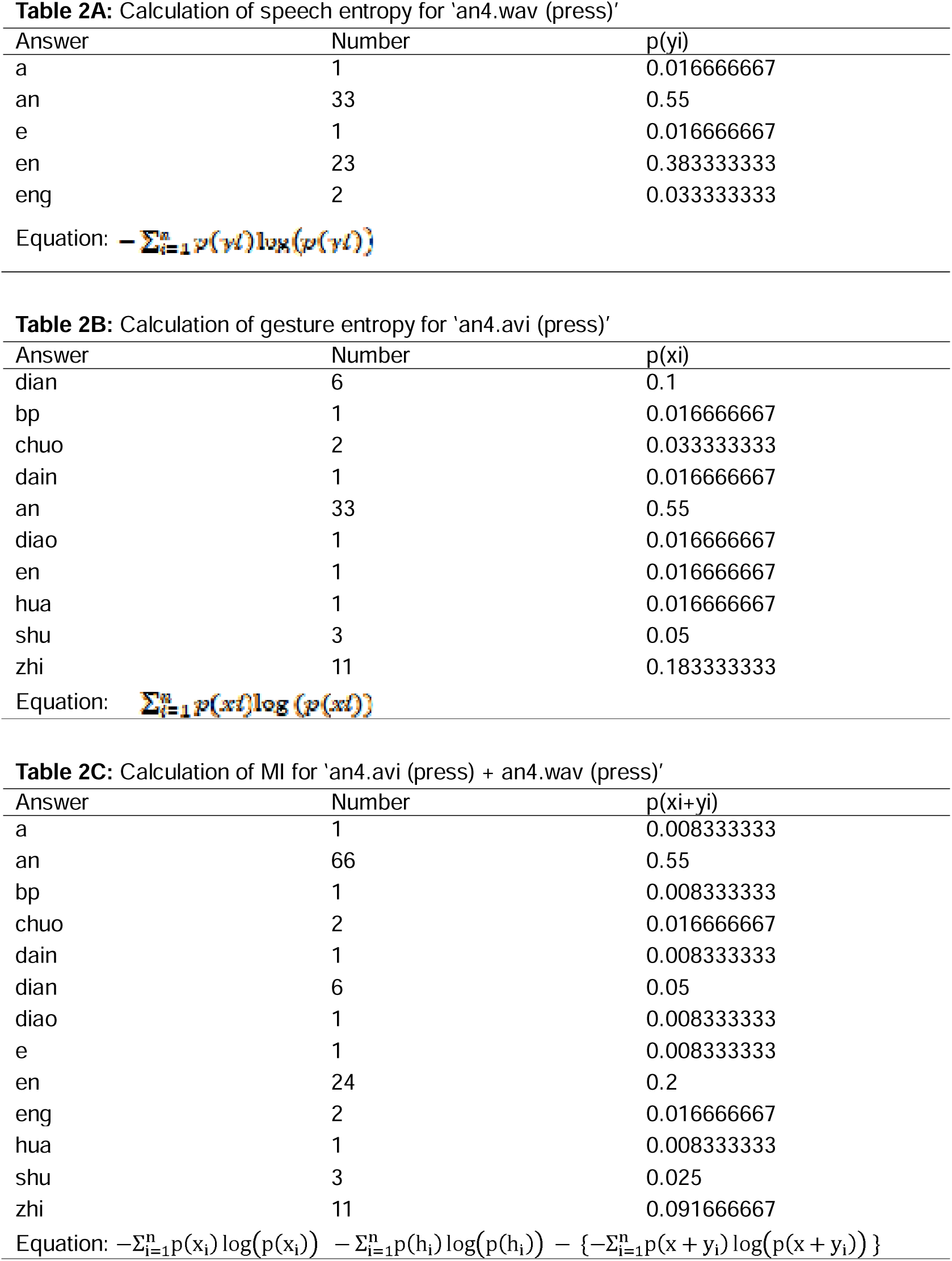
Examples of ‘an4 (press)’ for the calculation of speech entropy, gesture entropy and mutual information (MI).

| Answer | Number | p(yi) |
| --- | --- | --- |
| a | 1 | 0.016666667 |
| an | 33 | 0.55 |
| e | 1 | 0.016666667 |
| en | 23 | 0.383333333 |
| eng | 2 | 0.033333333 |

| Answer | Number | p(xi) |
| --- | --- | --- |
| dian | 6 | 0.1 |
| bp | 1 | 0.016666667 |
| chuo | 2 | 0.033333333 |
| dain | 1 | 0.016666667 |
| an | 33 | 0.55 |
| diao | 1 | 0.016666667 |
| en | 1 | 0.016666667 |
| hua | 1 | 0.016666667 |
| shu | 3 | 0.05 |
| zhi | 11 | 0.183333333 |

| Answer | Number | p(xi+yi) |
| --- | --- | --- |
| a | 1 | 0.008333333 |
| an | 66 | 0.55 |
| bp | 1 | 0.008333333 |
| chuo | 2 | 0.016666667 |
| dain | 1 | 0.008333333 |
| dian | 6 | 0.05 |
| diao | 1 | 0.008333333 |
| e | 1 | 0.008333333 |
| en | 24 | 0.2 |
| eng | 2 | 0.016666667 |
| hua | 1 | 0.008333333 |
| shu | 3 | 0.025 |
| zhi | 11 | 0.091666667 |

Equation: $-\sum_{i=1}^n p(x_i) \log(p(x_i)) - \sum_{i=1}^n p(h_i) \log(p(h_i)) - \{-\sum_{i=1}^n p(x + y_i) \log(p(x + y_i))\}$

**Supplementary Table 3.** Quantitative information for each stimulus.

| Stimuli<sup>1325</sup> | Gesture entropy | Speech entropy | Joint entropy | Mutual information | Gesture response No. | Speech response No. | Total response No. |
| --- | --- | --- | --- | --- | --- | --- | --- |
| an4 (press) | 2.13 | 1.37 | 2.15 | 1.35 | 10 | 5 | 13 |
| bai1 (break) | 0.91 | 0.11 | 0.61 | 0.41 | 5 | 2 | 6 |
| ca1 (wipe) | 2.07 | 0.56 | 1.67 | 0.96 | 10 | 3 | 10 |
| ding1 (hammer) | 2.55 | 0.00 | 1.28 | 1.27 | 11 | 1 | 11 |
| feng2 (sew) | 3.04 | 0.00 | 1.95 | 1.09 | 14 | 1 | 14 |
| ji3 (squeeze) | 2.50 | 0.00 | 1.86 | 0.64 | 11 | 1 | 11 |
| jian3 (cut) | 0.12 | 0.00 | 0.07 | 0.05 | 2 | 1 | 2 |
| jiao3 (stir) | 1.83 | 0.63 | 1.46 | 1.01 | 10 | 3 | 12 |
| ju4 (saw) | 4.34 | 0.00 | 2.77 | 1.57 | 25 | 1 | 25 |
| ning3 (twist) | 0.59 | 0.63 | 0.53 | 0.69 | 5 | 3 | 7 |
| pen1 (spray) | 3.29 | 0.80 | 2.61 | 1.49 | 17 | 4 | 20 |
| qie1 (slice) | 3.23 | 0.47 | 2.31 | 1.38 | 18 | 2 | 19 |
| reng1 (throw) | 1.59 | 0.29 | 1.09 | 0.79 | 9 | 2 | 9 |
| sao1 (sweep) | 4.01 | 1.12 | 4.47 | 0.66 | 23 | 4 | 18 |
| shan1 (slap) | 1.54 | 0.33 | 1.10 | 0.78 | 10 | 3 | 12 |
| si1 (tear) | 0.21 | 0.00 | 0.12 | 0.09 | 2 | 1 | 2 |
| ti2 (lift) | 1.48 | 0.00 | 0.88 | 0.60 | 9 | 1 | 9 |
| tui1 (push) | 1.62 | 0.00 | 0.96 | 0.66 | 10 | 1 | 17 |
| yao2 (shake) | 4.26 | 0.12 | 2.93 | 1.46 | 25 | 2 | 26 |
| yun4 (iron) | 4.15 | 0.00 | 2.78 | 1.37 | 25 | 1 | 25 |

**Supplementary Table 4.** RT of Sc and Si in three HD-tDCS stimulation conditions for IFG and pMTG.

|  | **Anodal** |  | **Cathodal** |  | **Sham** |  |
| --- | --- | --- | --- | --- | --- | --- |
|  | Sc (ms)<br>(Rt±SE) | Si (ms)<br>(Rt±SE) | Sc (ms)<br>(Rt±SE) | Si(ms)<br>(Rt±SE) | Sc (ms)<br>(Rt±SE) | Si(ms)<br>(Rt±SE) |
| **tDCS over IFG** | 521.95<br>±13.41 | 537.46<br>±15.05 | 518.41<br>±11.95 | 530.33<br>±13.01 | 513.96<br>±14.40 | 537.46<br>±15.53 |
| **tDCS over pMTG** | 531.94<br>±11.43 | 553.61<br>±13.43 | 531.88<br>±11.43 | 545.08<br>±11.97 | 545.08<br>±11.97 | 569.57<br>±14.32 |

**Supplementary figure 1.**
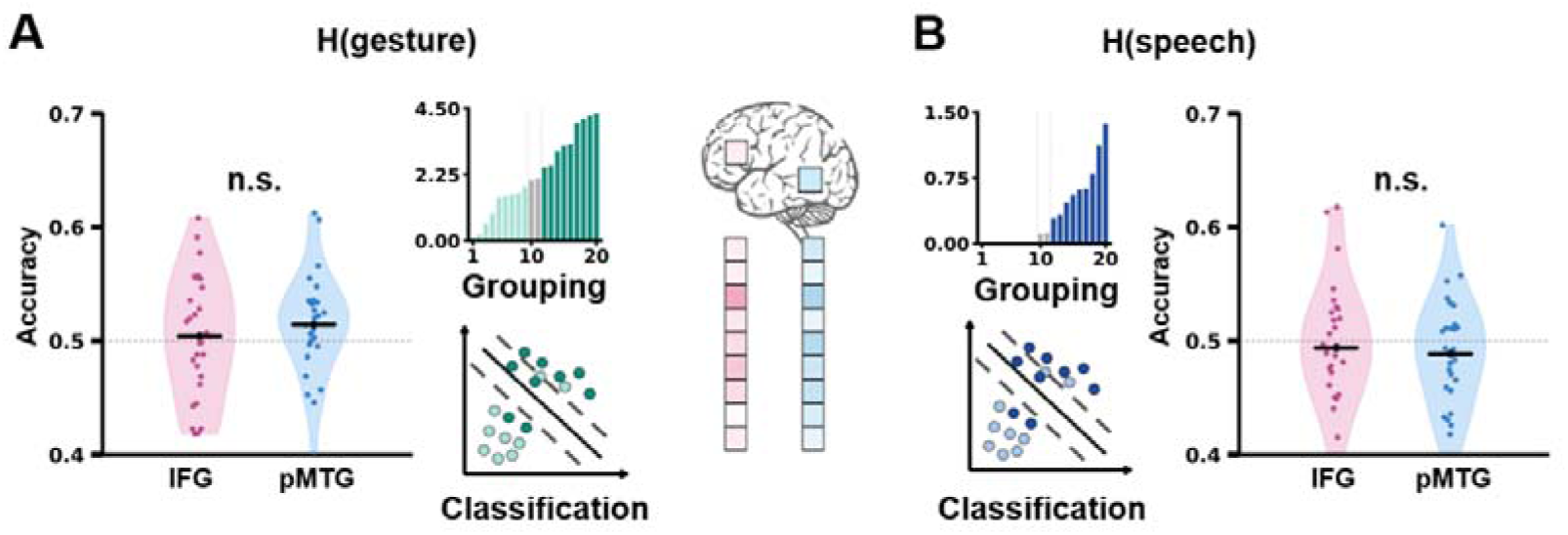
fMRI decoding of unimodal entropy within the IFG–pMTG network. **(A)** ROI-based multivoxel pattern analysis (MVPA) of gesture entropy. Stimuli were divided into low- and high-gesture-entropy groups, and linear support vector machine (SVM) classifiers were trained on beta-series multivoxel patterns within the predefined IFG and pMTG regions. Decoding accuracy did not significantly exceed chance in either region. **(B)** ROI-based MVPA of speech entropy. Stimuli were divided into low- and high-speech-entropy groups and classified using the same procedure. Decoding accuracy did not significantly exceed chance in either IFG or pMTG. Violin plots show participant-level balanced decoding accuracies; dots represent individual participants and horizontal black lines indicate group means. The dashed horizontal line indicates chance-level performance (0.50). n.s., not significant.

**Supplementary figure 2.**
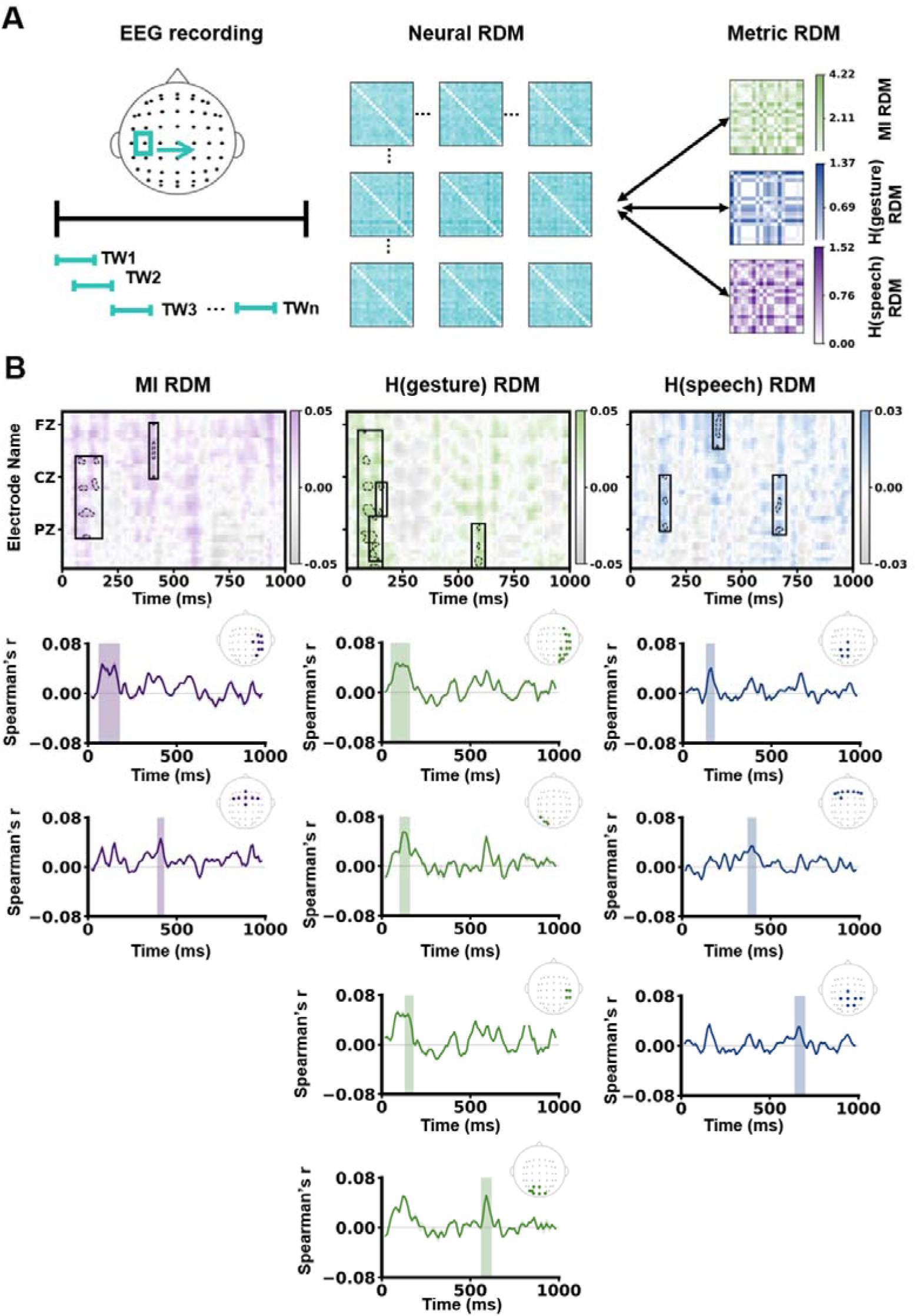
Whole-scalp time-resolved RSA of information-theoretic representations. **(A)** Schematic illustration of the whole-scalp representational similarity analysis (RSA). Neural representational dissimilarity matrices (RDMs) were constructed for each electrode across sliding time windows and compared with model RDMs derived from mutual information (MI), gesture entropy, and speech entropy using Spearman correlation. **(B)** Whole-scalp time-resolved RSA results. **Top**, spatiotemporal maps of RSA coefficients across electrodes and time; outlined regions indicate significant clusters identified by cluster-based permutation testing. **Bottom**, time courses of RSA coefficients for each significant cluster, with corresponding scalp distributions. Shaded regions indicate the temporal extent of significant clusters. All clusters were identified using spatiotemporal cluster-based permutation tests (*p* < 0.05).

## Methods and Results

To determine whether information-related representational structures extended beyond the predefined IFG- and pMTG-approximating electrode clusters, we conducted a complementary whole-scalp time-resolved representational similarity analysis (RSA).

For each participant, electrode, and time window, stimulus-specific neural responses were extracted from the ERP data, and neural representational dissimilarity matrices (RDMs) were constructed by calculating pairwise Euclidean distances among the 20 stimuli. Model RDMs were constructed separately for gesture entropy, speech entropy, and mutual information (MI) from pairwise Euclidean distances between their stimulus-level values. Neural and model RDMs were compared using Spearman correlations between their upper-triangular elements, and correlation coefficients were Fisher z-transformed before group-level statistical analysis.

At each electrode and time window, RSA effects were tested against zero using one-sample t-tests. Multiple comparisons across electrodes and time were controlled using spatiotemporal cluster-based permutation tests with sign-flipping across participants (5,000 permutations). Clusters were considered significant at a cluster-level corrected threshold of p < 0.05.

Because approximately half of the stimuli had speech entropy values of zero, the continuous speech-entropy model contained limited variation at the lower end of the distribution. We therefore performed an additional analysis using a binary speech-information model (0 = absent, 1 = present), with the corresponding model RDM representing whether stimulus pairs differed in the presence or absence of speech information. The same RSA and cluster-based permutation procedures were applied to this binary model.

Whole-scalp RSA revealed significant positive spatiotemporal clusters for MI at three processing stages. An early cluster was observed at 60–180 ms over right central–parietal electrodes (FC4, FC6, C2, C4, C6, CP4, CP6, and P4; mean *r* = 0.0513, mean *t* = 2.399, *p* = 0.002). A second cluster emerged at 390–430 ms over frontocentral electrodes (Fz, FC3, FC1, FCz, FC2, FC4, and Cz; mean *r* = 0.0465, mean *t* = 2.348, *p* = 0.020) (**Fig. S2 left**).

Gesture entropy also showed significant positive representational correspondence across multiple time windows, including clusters at 50–160 ms; mean *r* = 0.0512, mean *t* = 2.569, *p* = 0.004), 100–160 ms (mean *r* = 0.0588, mean *t* = 2.225, *p* = 0.022), 130–180 ms (mean *r* = 0.0491, mean *t* = 2.293, *p* = 0.029), and 560–620 ms (mean *r* = 0.0516, mean *t* = 2.311, *p* = 0.027) (**Fig. S2 middle**).

No significant positive spatiotemporal clusters were identified for the continuous speech-entropy model. The complementary binary speech-information model, however, revealed three significant positive clusters: 130–180 ms (mean *r* = 0.0407, mean *t* = 2.231, *p* = 0.011), 370–420 ms (mean *r* = 0.0347, mean *t* = 2.419, *p* = 0.010), and 640–700 ms (mean *r* = 0.0321, mean *t* = 2.320, *p* = 0.022) (**Fig. S2 right**).

Thus, whole-scalp RSA revealed temporally distributed representational effects for both modality-specific entropy and cross-modal MI. MI-related representational geometry shifted from right central– parietal electrodes during early processing to frontocentral electrodes during the N400-associated period and left frontal electrodes at later stages. Notably, the intermediate MI cluster overlapped temporally and topographically with the IFG-approximating RSA effect identified in the predefined electrode-cluster analysis (390-430ms), providing convergent evidence for MI-related representational structure during the semantic integration period.

**Supplementary figure 3.**
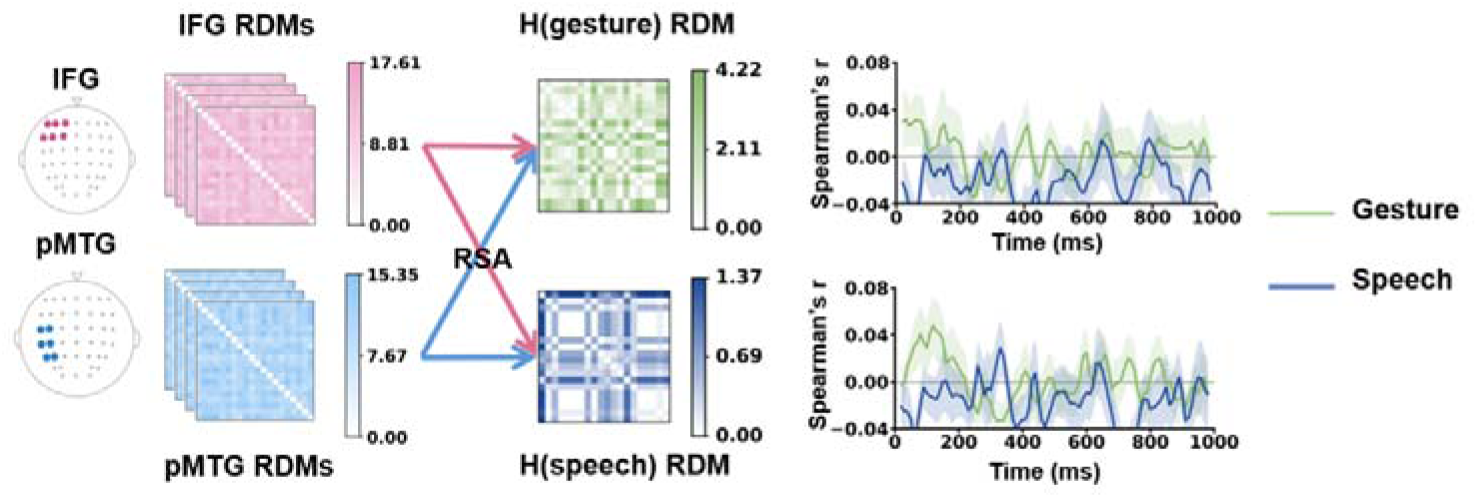
ROI-based time-resolved RSA of modality-specific entropy. Time-resolved representational similarity analysis (RSA) within predefined IFG- and pMTG-approximating electrode clusters. Neural representational dissimilarity matrices (RDMs) were constructed from multichannel EEG patterns across sliding time windows and compared with model RDMs derived from gesture entropy and speech entropy using Spearman correlation. Time courses show group-level RSA coefficients for gesture entropy (green) and speech entropy (blue) within the IFG (top) and pMTG (bottom) electrode clusters; shaded areas indicate variability across participants. No significant temporal clusters were identified for either entropy measure after cluster-based permutation correction.

**Supplementary figure 4.**
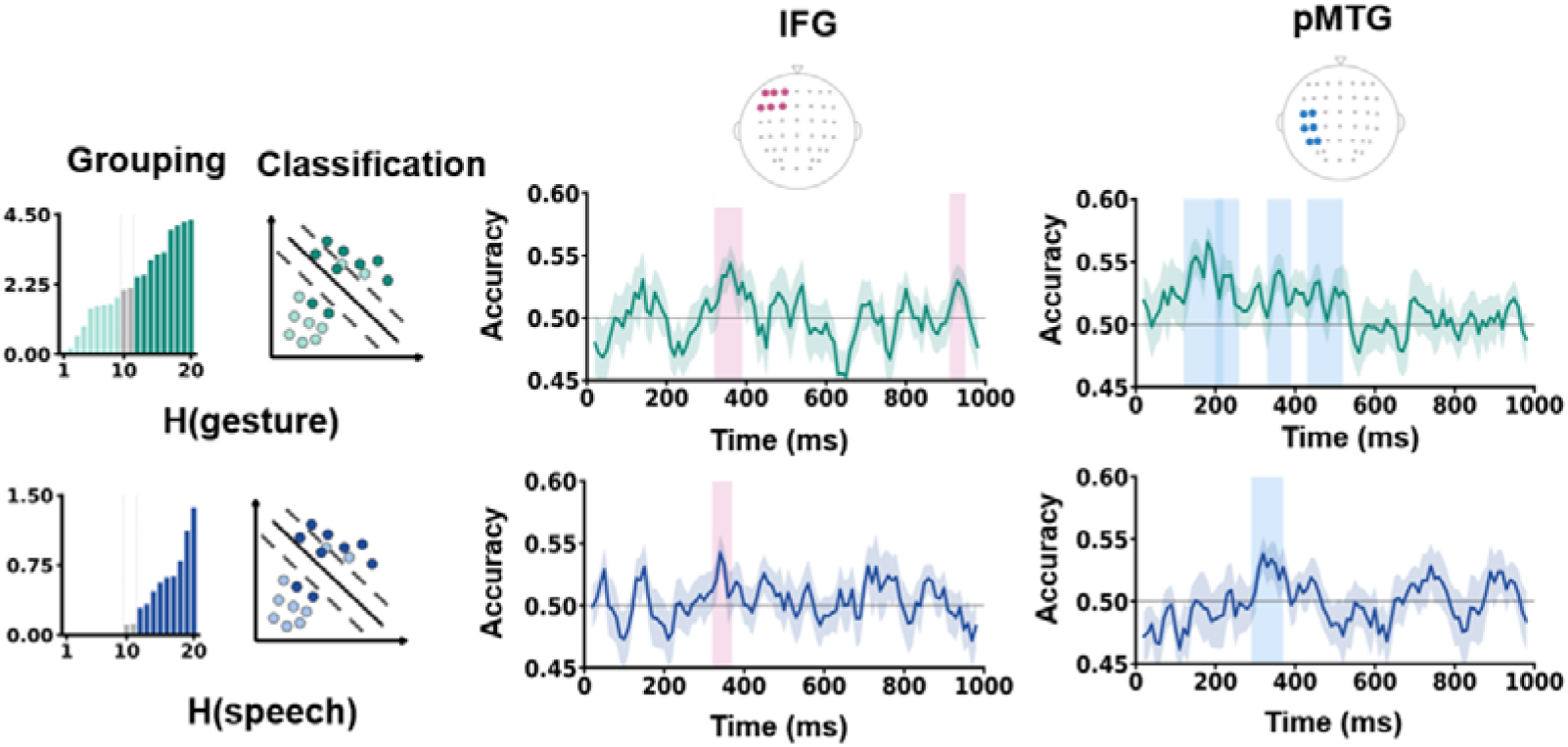
ROI-based time-resolved decoding of modality-specific entropy. Time-resolved multivariate decoding of gesture and speech entropy within predefined IFG- and pMTG-approximating electrode clusters. Stimuli were divided into low- and high-entropy groups, and linear support vector machine (SVM) classifiers were trained to discriminate the two conditions from multichannel EEG patterns across sliding time windows. Left, schematic illustration of the binary grouping and classification procedure. Middle and right, time-resolved decoding accuracy for gesture entropy (top) and speech entropy (bottom) within the IFG and pMTG electrode clusters, respectively.

For gesture entropy, significant above-chance decoding was observed in the IFG at 320– 390 ms (mean accuracy = 0.536, *p* = 0.012) and 910–950 ms (mean accuracy = 0.530, *p* = 0.047), and in the pMTG at 120–220 ms (mean accuracy = 0.550, *p* = 0.001), 200–260 ms (mean accuracy = 0.539, *p* = 0.011), 330–390 ms (mean accuracy = 0.539, *p* = 0.011), 430–480 ms (mean accuracy = 0.534, *p* = 0.024), and 480–520 ms (mean accuracy = 0.530, *p* = 0.050).

For speech entropy, significant decoding was observed at 320–370 ms in the IFG (mean accuracy = 0.538, *p* = 0.008) and 290–370 ms in the pMTG (mean accuracy = 0.532, *p* = 0.010).

Colored vertical bands indicate significant temporal clusters identified by cluster-based permutation testing; shaded areas around the decoding curves indicate variability across participants. The horizontal line indicates chance-level accuracy (0.50).

**Supplementary figure 5.**
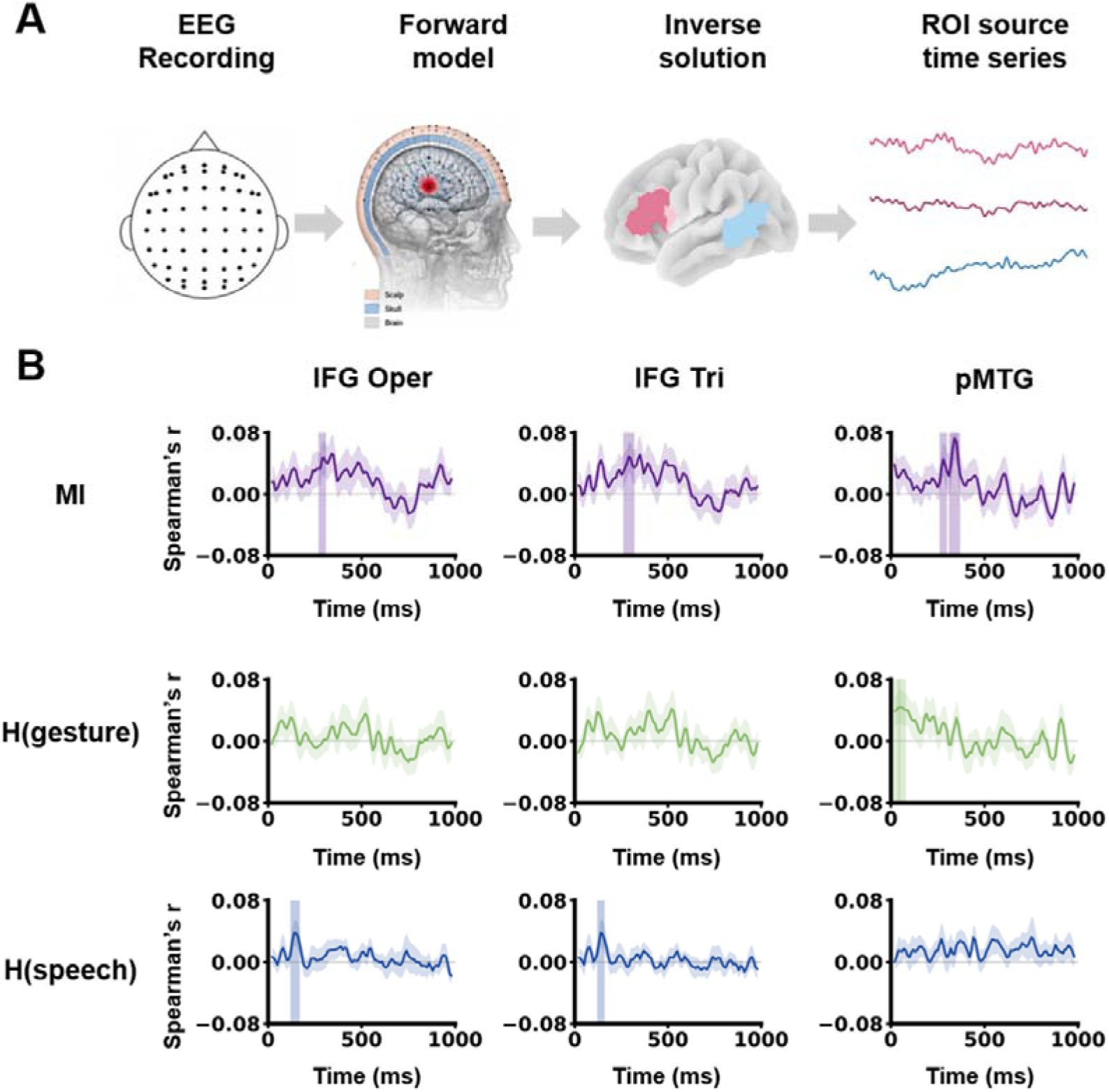
Source-space time-resolved RSA of information-theoretic representations. **(A)** Schematic illustration of the source-space representational similarity analysis (RSA). EEG signals were reconstructed in source space using forward modeling and inverse estimation, and source-level time series were extracted from predefined regions of interest: the left inferior frontal gyrus pars opercularis (IFG oper), pars triangularis (IFG tri), and posterior middle temporal gyrus (pMTG). Neural representational dissimilarity matrices (RDMs) were constructed across sliding time windows and compared with model RDMs derived from mutual information (MI), gesture entropy, and speech entropy using Spearman correlation. **(B)** Time-resolved source-space RSA within the three ROIs. Rows show RSA time courses for MI (**top**), gesture entropy (**middle**), and speech entropy (**bottom**); columns show IFG oper, IFG tri, and pMTG, respectively. Curves indicate group-level Spearman correlation coefficients between neural and model RDMs across time, with surrounding bands indicating variability across participants. Colored vertical bands indicate significant temporal clusters identified by cluster-based permutation testing (*p* < 0.05).

## Method and results

To provide an anatomical complement to the scalp-level representational effects, source-space RSA was performed to examine whether information-theoretic representations were expressed within cortical regions implicated in semantic processing.

Source reconstruction was performed using a standard cortical source model based on the fsaverage template and a distributed inverse solution implemented using standardized low-resolution electromagnetic tomography (sLORETA). Sensor-level EEG activity was projected into cortical source space to estimate stimulus-locked source activity. Source-space analyses focused on three anatomically defined regions of interest (ROIs): the left inferior frontal gyrus pars opercularis (IFG oper), left inferior frontal gyrus pars triangularis (IFG tri), and left posterior middle temporal gyrus (pMTG).

Anatomical ROIs were defined according to the Automated Anatomical Labeling atlas (AAL3). The pMTG ROI was defined as the posterior half of the left middle temporal gyrus, separated at the median MNI y-coordinate of all voxels within the anatomical region, providing a closer approximation to the posterior temporal region examined in the fMRI analyses.

For each ROI and time window, neural representational dissimilarity matrices (RDMs) were constructed from pairwise Euclidean distances in ROI-averaged source activity across the 20 stimuli. Model RDMs for gesture entropy, speech entropy, mutual information (MI), and the complementary binary speech-information model were constructed using the same procedure as in the scalp-level RSA. Neural and model RDMs were compared using Spearman correlations between their upper-triangular elements, and correlation coefficients were Fisher z-transformed before group-level analysis. RSA effects were tested against zero using one-sample t-tests, with temporal cluster-based permutation testing based on sign-flipping across participants (5,000 permutations; cluster-level corrected *p* < 0.05).

Source-space RSA revealed significant MI-related representational correspondence across all three predefined ROIs. In the left IFG pars opercularis, MI-related representational similarity was significant at 270–310 ms (mean *r* = 0.0472, mean *t* = 2.256, *p* = 0.040), while the pars triangularis showed a significant effect at 260–320 ms (mean *r* = 0.0456, mean *t* = 2.295, *p* = 0.029). In the pMTG, significant MI-related clusters were identified at 260–300 ms (mean *r* = 0.0458, mean *t* = 2.164, *p* = 0.033) and 310–370 ms (mean *r* = 0.0681, mean *t* = 2.390, *p* = 0.029) (**Fig. S5 B top**). Thus, MI-related representational effects converged across frontal and posterior temporal source ROIs within an approximately 260–370 ms interval.

Gesture entropy showed a more restricted source-space pattern, with significant representational correspondence observed only in the pMTG during two early temporal clusters (10–50 ms and 40–80 ms). No significant positive clusters were identified for gesture entropy in either IFG ROI (**Fig. S5 B middle**).

Similarly, the continuous speech-entropy model showed no significant positive source-space clusters. The complementary binary speech-information model, however, revealed significant effects in the IFG pars opercularis at 120–170 ms and in the pars triangularis at 120–160 ms (**Fig. S5 B bottom**).

Overall, the source-space analysis complemented the sensor-space RSA by revealing MI-related representational structure across both IFG and pMTG source ROIs during the 260–370 ms period. These source-level effects occurred slightly earlier than, but temporally adjacent to, the 390–430 ms frontocentral MI-related effect identified in the sensor-space analysis, providing convergent evidence for MI-related representational dynamics around the N400-associated semantic integration period.

## References

1. Damasio, H., Grabowski, T.J., Tranel, D., Hichwa, R.D., and Damasio, A.R. (1996). A neural basis for lexical retrieval. Nature 380, 499–505. DOI 10.1038/380499a0.

2. Patterson, K., Nestor, P.J., and Rogers, T.T. (2007). Where do you know what you know? The representation of semantic knowledge in the human brain. Nature Reviews Neuroscience 8, 976–987. 10.1038/nrn2277.

3. Brennan, J.R., Stabler, E.P., Van Wagenen, S.E., Luh, W.M., and Hale, J.T. (2016). Abstract linguistic structure correlates with temporal activity during naturalistic comprehension. Brain and Language 157, 81–94. 10.1016/j.bandl.2016.04.008.

4. Benetti, S., Ferrari, A., and Pavani, F. (2023). Multimodal processing in face-to-face interactions: A bridging link between psycholinguistics and sensory neuroscience. Front Hum Neurosci 17, 1108354. 10.3389/fnhum.2023.1108354.

5. Noppeney, U. (2021). Perceptual Inference, Learning, and Attention in a Multisensory World. Annual Review of Neuroscience, Vol 44, 2021 44, 449-473. 10.1146/annurev-neuro-100120-085519.

6. Ma, W.J., and Jazayeri, M. (2014). Neural coding of uncertainty and probability. Annu Rev Neurosci 37, 205–220. 10.1146/annurev-neuro-071013-014017.

7. Fischer, B.J., and Pena, J.L. (2011). Owl’s behavior and neural representation predicted by Bayesian inference. Nat Neurosci 14, 1061–1066. 10.1038/nn.2872.

8. Ganguli, D., and Simoncelli, E.P. (2014). Efficient sensory encoding and Bayesian inference with heterogeneous neural populations. Neural Comput 26, 2103–2134. 10.1162/NECO_a_00638.

9. Hostetter, A., and Mainela-Arnold, E. (2015). Gestures occur with spatial and Motoric knowledge: It’s more than just coincidence. Perspectives on Language Learning and Education 22, 42–49. doi:10.1044/lle22.2.42.

10. McNeill, D. (2005). Gesture and though (University of Chicago Press). 10.7208/chicago/9780226514642.001.0001.

11. Kendon, A. (1997). Gesture. Annu Rev Anthropol 26, 109–128. 10.1146/annurev.anthro.26.1.109.

12. Hagoort, P. (2005). On broca, brain, and binding: a new framework. Trends in Cognitive Sciences 9, 416–423. 10.1016/j.tics.2005.07.004.

13. Hagoort, P., Hald, L., Bastiaansen, M., and Petersson, K.M. (2004). Integration of word meaning and world knowledge in language comprehension. Science 304, 438–441. 10.1126/science.1095455.

14. Ozyurek, A., Willems, R.M., Kita, S., and Hagoort, P. (2007). On-line integration of semantic information from speech and gesture: Insights from event-related brain potentials. J Cognitive Neurosci 19, 605–616. 10.1162/jocn.2007.19.4.605.

15. Willems, R.M., Ozyurek, A., and Hagoort, P. (2009). Differential roles for left inferior frontal and superior temporal cortex in multimodal integration of action and language. Neuroimage 47, 1992–2004. 10.1016/j.neuroimage.2009.05.066.

16. Drijvers, L., Jensen, O., and Spaak, E. (2021). Rapid invisible frequency tagging reveals nonlinear integration of auditory and visual information. Human Brain Mapping 42, 1138–1152. 10.1002/hbm.25282.

17. Drijvers, L., and Ozyurek, A. (2018). Native language status of the listener modulates the neural integration of speech and iconic gestures in clear and adverse listening conditions. Brain and Language 177, 7–17. 10.1016/j.bandl.2018.01.003.

18. Drijvers, L., van der Plas, M., Ozyurek, A., and Jensen, O. (2019). Native and non-native listeners show similar yet distinct oscillatory dynamics when using gestures to access speech in noise. Neuroimage 194, 55–67. 10.1016/j.neuroimage.2019.03.032.

19. Holle, H., and Gunter, T.C. (2007). The role of iconic gestures in speech disambiguation: ERP evidence. J Cognitive Neurosci 19, 1175–1192. 10.1162/jocn.2007.19.7.1175.

20. Kita, S., and Ozyurek, A. (2003). What does cross-linguistic variation in semantic coordination of speech and gesture reveal?: Evidence for an interface representation of spatial thinking and speaking. J Mem Lang 48, 16–32. 10.1016/S0749-596x(02)00505-3.

21. Bernardis, P., and Gentilucci, M. (2006). Speech and gesture share the same communication system. Neuropsychologia 44, 178–190. 10.1016/j.neuropsychologia.2005.05.007.

22. Zhao, W.Y., Riggs, K., Schindler, I., and Holle, H. (2018). Transcranial magnetic stimulation over left inferior frontal and posterior temporal cortex disrupts gesture-speech integration. Journal of Neuroscience 38, 1891–1900. 10.1523/Jneurosci.1748-17.2017.

23. Zhao, W., Li, Y., and Du, Y. (2021). TMS reveals dynamic interaction between inferior frontal gyrus and posterior middle temporal gyrus in gesture-speech semantic integration. The Journal of Neuroscience, 10356–10364. 10.1523/jneurosci.1355-21.2021.

24. Shannon, C.E. (1948). A mathematical theory of communication. Bell Syst Tech J 27, 379–423. 10.1002/j.1538-7305.1948.tb01338.x.

25. Tremblay, P., Deschamps, I., Baroni, M., and Hasson, U. (2016). Neural sensitivity to syllable frequency and mutual information in speech perception and production. Neuroimage 136, 106–121. 10.1016/j.neuroimage.2016.05.018.

26. Bikson, M., Inoue, M., Akiyama, H., Deans, J.K., Fox, J.E., Miyakawa, H., and Jefferys, J.G.R. (2004). Effects of uniform extracellular DC electric fields on excitability in rat hippocampal slices. J Physiol-London 557, 175–190. 10.1113/jphysiol.2003.055772.

27. Federmeier, K.D., Mai, H., and Kutas, M. (2005). Both sides get the point: hemispheric sensitivities to sentential constraint. Memory & Cognition 33, 871–886. 10.3758/bf03193082.

28. Kelly, S.D., Kravitz, C., and Hopkins, M. (2004). Neural correlates of bimodal speech and gesture comprehension. Brain and Language 89, 253–260. 10.1016/s0093-934x(03)00335-3.

29. Wu, Y.C., and Coulson, S. (2005). Meaningful gestures: Electrophysiological indices of iconic gesture comprehension. Psychophysiology 42, 654–667. 10.1111/j.1469-8986.2005.00356.x.

30. Fritz, I., Kita, S., Littlemore, J., and Krott, A. (2021). Multimodal language processing: How preceding discourse constrains gesture interpretation and affects gesture integration when gestures do not synchronise with semantic affiliates. J Mem Lang 117, 104191. 10.1016/j.jml.2020.104191.

31. Gunter, T.C., and Weinbrenner, J.E.D. (2017). When to take a gesture seriously: On how we use and prioritize communicative cues. J Cognitive Neurosci 29, 1355–1367. 10.1162/jocn_a_01125.

32. Rogers, T.T., Ralph, M.A.L., Garrard, P., Bozeat, S., McClelland, J.L., Hodges, J.R., and Patterson, K. (2004). Structure and deterioration of semantic memory: A neuropsychological and computational investigation. Psychological Review 111, 205–235. 10.1037/0033-295x.111.1.205.

33. Ralph, M.A.L., Jefferies, E., Patterson, K., and Rogers, T.T. (2017). The neural and computational bases of semantic cognition. Nature Reviews Neuroscience 18, 42–55. 10.1038/nrn.2016.150.

34. Rogers, T.T., Hodges, J.R., Ralph, M.A.L., and Patterson, K. (2003). Object recognition under semantic impairment: The effects of conceptual regularities on perceptual decisions. Language and Cognitive Processes 18, 625–662. 10.1080/01690960344000053.

35. Martin, S., Ferrante, M., Bruera, A., et al. (2025). Causal contributions of left inferior and medial frontal cortex to semantic and executive control. Communications Biology 8, 1343 10.1038/s42003-025-08848-5.

36. Jackson, R.L. (2021). The neural correlates of semantic control revisited. NeuroImage 224, 117444. 10.1016/j.neuroimage.2020.117444.

37. Fernandino, L., Binder, J.R., Desai, R.H., Pendl, S.L., Humphries, C.J., Gross, W.L., Conant, L.L., and Seidenberg, M.S. (2016). Concept representation reflects multimodal abstraction: A framework for embodied semantics. Cerebral Cortex 26, 2018–2034 10.1093/cercor/bhv020.

38. Fadiga, L., Craighero, L., and Olivier, E. (2005). Human motor cortex excitability during the perception of others’ action. Current Opinion in Neurobiology 15, 213–218. 10.1016/j.conb.2005.03.013.

39. Giard, M.H., and Peronnet, F. (1999). Auditory-visual integration during multimodal object recognition in humans: A behavioral and electrophysiological study. Journal of Cognitive Neuroscience 11, 473–490. 10.1162/089892999563544.

40. Meijer, G.T., Mertens, P.E.C., Pennartz, C.M.A., Olcese, U., and Lansink, C.S. (2019). The circuit architecture of cortical multisensory processing: Distinct functions jointly operating within a common anatomical network. Progress in Neurobiology 174, 1–15. 10.1016/j.pneurobio.2019.01.004.

41. Senkowski, D., and Engel, A.K. (2024). Multi-timescale neural dynamics for multisensory integration. Nature Reviews Neuroscience 25, 625–642. 10.1038/s41583-024-00845-7.

42. Bizley, J.K., Maddox, R.K., and Lee, A.K.C. (2016). Defining auditory-visual objects: Behavioral tests and physiological mechanisms. Trends in Neurosciences 39, 74–85. 10.1016/j.tins.2015.12.007.

43. Kutas, M., and Federmeier, K.D. (2011). Thirty years and counting: Finding meaning in the N400 component of the event-related brain potential (ERP). Annual Review of Psychology 62, 621–647. 10.1146/annurev.psych.093008.131123.

44. Tesink, C.M.J.Y., Petersson, K.M., van Berkum, J.J.A., van den Brink, D., Buitelaar, J.K., and Hagoort, P. (2009). Unification of speaker and meaning in language comprehension: An fMRI study. Journal of Cognitive Neuroscience 21, 2085–2099. 10.1162/jocn.2008.21161.

45. McNeill, D. (1992). Hand and Mind: What Gestures Reveal about Thought. University of Chicago Press. 10.2307/1576015.

46. Steuer, R., Kurths, J., Daub, C.O., Weise, J., and Selbig, J. (2002). The mutual information: Detecting and evaluating dependencies between variables. Bioinformatics 18, S231–S240. 10.1093/bioinformatics/18.

47. Matyjek, M., Kita, S., Torralba Cuello, M., and Soto-Faraco, S. (2024). Multisensory integration of speech and gestures in a naturalistic paradigm. Human Brain Mapping 45, e26797. 10.1002/hbm.26797.

48. Zhao, W. (2023). TMS reveals a two-stage priming circuit of gesture-speech integration. Frontiers in Psychology 14, 1156087. 10.3389/fpsyg.2023.1156087.

49. Kelly, S.D., Creigh, P., and Bartolotti, J. (2010). Integrating speech and iconic gestures in a Stroop-like task: Evidence for automatic processing. Journal of Cognitive Neuroscience 22, 683–694. 10.1162/jocn.2009.21254.

50. Koessler, L., Maillard, L., Benhadid, A., Vignal, J.P., Felblinger, J., Vespignani, H., and Braun, M. (2009). Automated cortical projection of EEG sensors: Anatomical correlation via the international 10-10 system. NeuroImage 46, 64–72. 10.1016/j.neuroimage.2009.02.006.

51. Mumford, J.A., Turner, B.O., Ashby, F.G., and Poldrack, R.A. (2012). Deconvolving BOLD activation in event-related designs for multivoxel pattern classification analyses. NeuroImage 59, 2636–2643. 10.1016/j.neuroimage.2011.08.076.

52. Pereira, F., Mitchell, T., and Botvinick, M. (2009). Machine learning classifiers and fMRI: A tutorial overview. NeuroImage 45, S199–S209. 10.1016/j.neuroimage.2008.11.007.

53. Delorme, A., and Makeig, S. (2004). EEGLAB: An open source toolbox for analysis of single-trial EEG dynamics including independent component analysis. Journal of Neuroscience Methods 134, 9–21. 10.1016/j.jneumeth.2003.10.009.

54. Habets, B., Kita, S., Shao, Z.S., Ozyurek, A., and Hagoort, P. (2011). The role of synchrony and ambiguity in speech-gesture integration during comprehension. Journal of Cognitive Neuroscience 23, 1845–1854. 10.1162/jocn.2010.21462.

55. Cichy, R.M., Pantazis, D., and Oliva, A. (2016). Similarity-based fusion of MEG and fMRI reveals spatio-temporal dynamics in human cortex during visual object recognition. Cerebral Cortex 26, 3563–3579. 10.1093/cercor/bhw135.

